# Spiral ganglion neuron degeneration in deafened rats involves innate and adaptive immune responses not requiring complement

**DOI:** 10.1101/2024.02.21.581500

**Authors:** Benjamin M. Gansemer, Muhammad T. Rahman, Zhenshen Zhang, Steven H. Green

## Abstract

Spiral ganglion neurons (SGNs) transmit auditory information from cochlear hair cells to the brain. SGNs are thus not only important for normal hearing, but also for effective functioning of cochlear implants, which stimulate SGNs when hair cells are missing. SGNs slowly degenerate following aminoglycoside-induced hair cell loss, a process thought to involve an immune response. However, the specific immune response pathways involved remain unknown. We used RNAseq to gain a deeper understanding immune-related and other transcriptomic changes that occur in the rat spiral ganglion after kanamycin-induced deafening. Among the immune and inflammatory genes that were selectively upregulated in deafened spiral ganglia, the complement cascade genes were prominent. We then assessed SGN survival, as well as immune cell infiltration and activation, in the spiral ganglia of rats with a CRISPR-Cas9-mediated knockout of complement component 3 (C3). Similar to previous findings in our lab and other deafened rodent models, we observed infiltration of macrophages and increased expression of CD68, a marker of phagocytic activity and cell activation, in the deafened ganglia. Moreover, we found that the immune response also includes MHCII+ macrophages and CD45+ and lymphocytes, indicative of an adaptive response. However, C3 knockout did affect SGN survival or macrophage infiltration/activation, indicating that complement activation does not play a role in SGN death after deafening. Together, these data suggest that both innate and adaptive immune responses are activated in the deafened spiral ganglion, with the adaptive response directly contributing to cochlear neurodegeneration.

## Introduction

Cochlear trauma leading to loss of hair cells can result in subsequent death and degeneration of spiral ganglion neurons (SGNs), the neurons that transmit auditory information from cochlear hair cells to the brainstem. In particular, hair cell loss caused by aminoglycoside exposure is known to induce SGN degeneration in rats (Alam et al., 2007), cats (Leake and Hradek, 1988; Leake et al., 2011), mice (Jansen et al., 2013), and humans (Jiang et al., 2017). The reasons for this cochlear neurodegeneration remain unclear, but accumulating evidence implicates an immune response. Macrophages which have a central role in the innate immune response, have been shown to be recruited to the spiral ganglion after trauma, augmenting the resident population. The increase in the number of macrophages in the spiral ganglion is accompanied by macrophage activation. These events have been shown in the ganglion after aminoglycoside-induced deafening in rats (Rahman et al., 2023) and mice (Kaur et al., 2018) and after noise-induced trauma in mice (Hirose et al., 2005; Kaur et al., 2018). Macrophages are early responders to damage or infection and are typically associated with protection. Consistent with this, some studies have shown macrophages playing a protective role in response to cochlear damage (Kaur et al., 2015; Kaur et al., 2018). In contrast, we have previously shown that anti-inflammatory drugs reduce the extent of SGN death (Rahman et al., 2023), suggesting a detrimental role for macrophages. It is plausible that there may be different populations of macrophages performing different functions (Bedeir et al., 2022). Additionally, immune cells typically associated with adaptive responses, including T cells and B cells, have been observed in the cochlea after noise-induced damage (Rai et al., 2020). However, the specific roles of lymphocytes, as well as macrophages, and the specific signals recruiting and activating immune cells in the cochlea, have yet to be uncovered.

Macrophages and other immune cells can be activated via the complement pathway. The complement pathway is generally thought of as part of the innate immune system, used for identification and clearance of dead cells and pathogens (Noris and Remuzzi, 2013). Complement can also assist in antibody-mediated immune functions, thereby contributing to adaptive immune responses (Lo and Woodruff, 2020). Complement has been shown to have beneficial effects, particularly related to synapse pruning, during brain development (Schafer et al., 2012). However, complement activation has also been implicated in the pathogenesis of many neurodegenerative diseases, including Alzheimer’s disease (Fonseca et al., 2004) and multiple sclerosis (Morgan et al., 2020), as well as peripheral neuropathies (Xu et al., 2018; Royer et al., 2019). Complement component 3 (C3) is involved in all pathways of complement activation and has been extensively studied in the context of CNS neurodegenerative diseases. C3 knockout or blockade has been shown to reduce neuroinflammation and improve neuronal function and cell survival in animal models of several neurodegenerative diseases (Schartz and Tenner, 2020).

Here we used RNAseq to further clarify the transcriptomic changes that occur in the spiral ganglion of aminoglycoside-deafened rats. We identified upregulation of the early complement components and receptors, among several immune response-related groups of genes. We further aimed to identify immune cell types present in the spiral ganglion during the period of SGN death and to clarify the role of complement in SGN degeneration post-deafening. We identified CD45+ lymphocytes in the hearing ganglion, with the number of CD45+ lymphocytes and of MHCII+ macrophages in the ganglion increasing after deafening. These results indicate that both innate and adaptive responses are active in the spiral ganglion after deafening, although each may contribute to different outcomes. However, we found that neither SGN survival nor macrophage infiltration and activation after deafening were affected by lack of C3.

## Materials and Methods

### Animals, deafening, and verification

For the RNAseq experiments, all animals used were Sprague-Dawley rats from our breeding colony or from dams purchased from Envigo. The C3 knockout (C3KO) rats, which have a CRISPR-generated insertion in the second exon resulting in early stop codon (Xu et al., 2018), were generously provided by Dr. Feng Lin (Cleveland Clinic). The C3KO rats, originally in an F344/Sprague-Dawley mixed background, were backcrossed into our Sprague-Dawley colony to generate C3-heterozygous animals. C3-heterozygous animals were mated to generate C3 wildtype (WT), C3 heterozygous, and C3KO littermates for deafening and histological analysis. Rat pups from the C3 heterozygous crosses were first genotyped by PCR by running two separate reactions for each animal: one to amplify the WT allele and one to amplify the mutant allele. The primer pairs used were: common forward primer: 5’-GTCTGCTCTGGCAGGTGTC-3‘ and reverse C3 WT-specific primer: 3‘-ACGAGTCCCACTACAGGGTC-5‘ or reverse C3 KO-specific primer: 3‘-GCAGGGTCCCACTACAGG-5‘. Genotypes were subsequently confirmed by Sanger sequencing using the common forward primer. Lack of C3 protein was additionally confirmed in a subset of rats using Western blot (Supplemental Figure 1D). The day of birth was designated as postnatal day 0 (P0). Both male and female rats were used in this study. Rats were housed under a 12 hr light/dark cycle and had free access to food and water. Protocols and procedures for all animal experiments were approved by the University of Iowa Institutional Animal Care and Use Committee.

Neonatal rats were deafened by daily intraperitoneal injections of kanamycin (400 mg/kg) from P8-16. Deafening was verified by lack of a detectable auditory brainstem response (ABR) to a 90 dB SPL click stimulus when tested between P24-P30 (Figure 1A). As previously reported, this method reliably eliminates inner and outer hair cells by P18 (Bailey and Green, 2014), and rats with no detectable ABR have no surviving hair cells (inner or outer) at P32 or P60 (Bailey and Green, 2014; Rahman et al., 2023). Any injected rats with a detectable ABR were excluded from the study.

**Figure 1.**
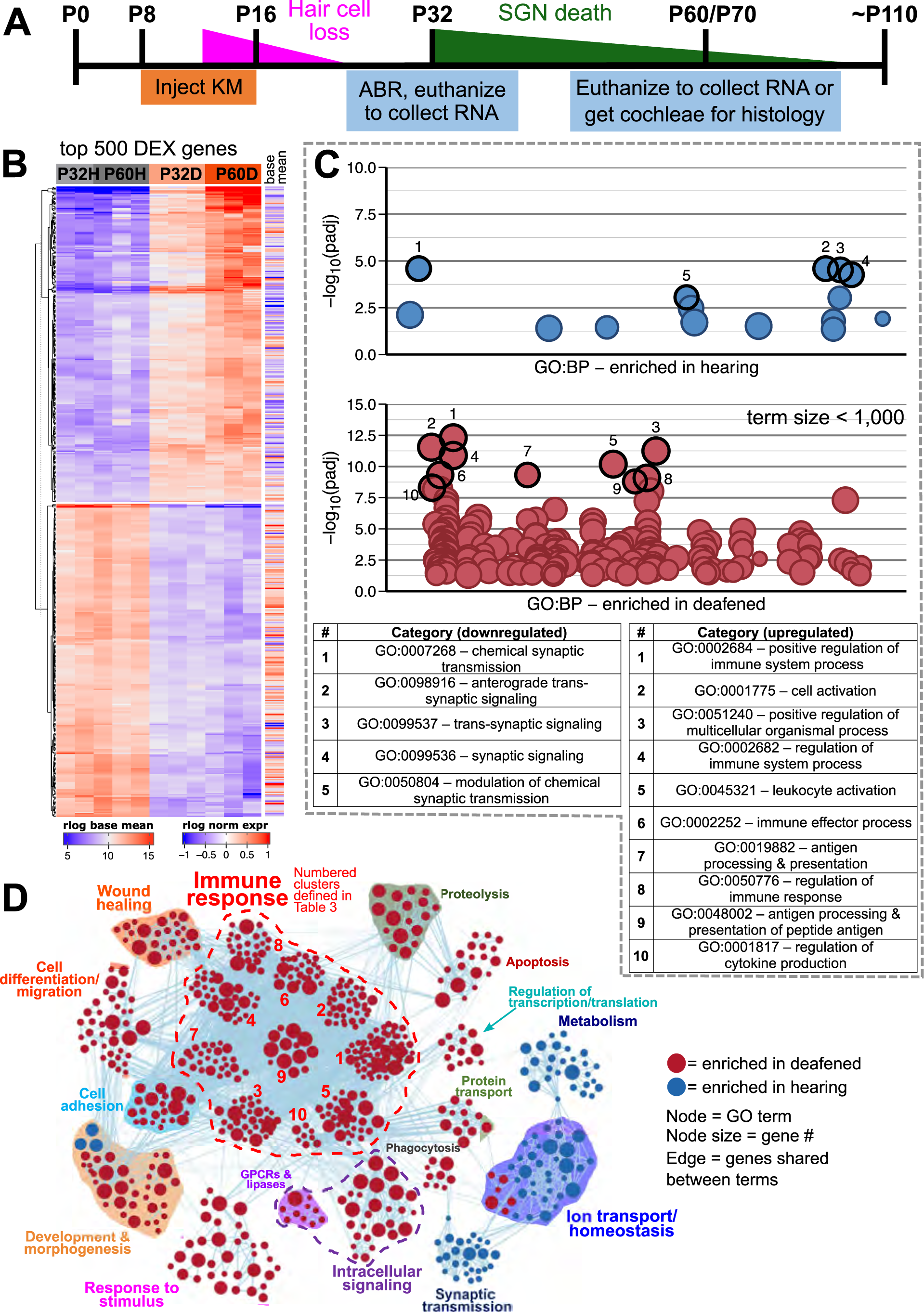
Transcriptome changes in the spiral ganglion after aminoglycoside-induced deafening. A) Timeline showing when rats were injected with kanamycin, the timing of hair cell loss and SGN death, and when cochleae were collected for RNA or histology. Cochleae were collected at P32 and P60 for RNA and at P70 for histology. B) Heatmap showing regularized logarithm (rlog) normalized expression of the top 500 differentially expressed genes (500 genes with the lowest padj). Base mean = average expression across all conditions/replicates. C) Manhattan plots showing gProfiler results when the genes in the heatmap in A were separated into downregulated (log_2_FC < 0, top plot) or upregulated (log_2_FC >0, bottom plot) and used as input. In the plot for upregulated genes, only terms with a size <1,000 are plotted. The top enriched categories in each plot are highlighted and the term names for each are shown in the table below the plots. D) GSEA network visualization showing enriched categories when comparing hearing vs. deaf, regardless of age (combined P32 and P60). Red nodes are GO categories enriched in deafened, blue nodes are those enriched in hearing. Edges (lines) indicate shared genes between the two connected terms. The category names for the numbered immune response clusters are defined in the tables below the network visualization – and are the same as used in Supplemental Figure 3A.

### Tissue recovery and preparation for RNA sequencing

The RNAseq experiment was performed with three biological replicates, with each biological replicate containing the spiral ganglia from 3–4 rats (6–8 ganglia). Ganglia from male and female rats were mixed so that each replicate contained tissue from both sexes. For tissue recovery, P32 or P60 rats were anesthetized using a ketamine (40 mg/kg)/xylazine (4 mg/kg) mixture and, while under surgical plane, killed by decapitation. The temporal bones were quickly isolated and the cochlea removed and placed in ice-cold phosphate buffered saline pH 7.4 (PBS) for dissection. The bony capsule was first removed, followed by the spiral ligament, stria vascularis, and cochlear epithelium/organ of Corti (Supplemental Figure 1A). The spiral ganglia were then placed in RNAlater (Thermofisher), flash-frozen in liquid nitrogen, and stored at −80°C until RNA extraction.

At the time of RNA extraction, when all samples had been collected, the samples in RNAlater were thawed and the RNAlater removed and replaced by Trizol (Invitrogen). The tissue was lysed and homogenized in Trizol using sterile stainless steel beads with the Qiagen TissueLyser LT (50Hz for 3 min). The lysate was then transferred to a pre-spun Phase Lock Gel Heavy (Quantabio, cat. #2302830) and the aqueous phase extracted using chloroform and centrifugation. RNA was purified from the aqueous phase using the Qiagen RNeasy mini kit (Qiagen) following manufacturer protocol. On-column DNase I treatment was performed following the Qiagen protocol. Purified RNA samples were flash-frozen in liquid nitrogen and stored at −80°C.

### Quantitative real-time PCR

For quantitative real-time polymerase chain reaction (qPCR), spiral ganglia were isolated at P60 as described above for RNA sequencing and placed in Trizol and stored at −80°C until RNA extraction. Ganglia from 3–4 rats were pooled for each biological replicate, keeping tissue from male and female rats separate. Three biological replicates from each of 4 conditions (hearing female, hearing male, deafened female, deafened male) were collected. RNA was extracted as described for RNA sequencing and cDNA was generated using the PrimeScript RT reagent kit (Takara, cat. #RR037A) using a mix of oligo dT and random hexamer primers. qPCR was done using PowerTrack SYBR Green Master Mix (Applied Biosystems, cat. #A46109) on an Applied Biosystems QuantStudio 3. Forward and reverse primers (Supplemental Table 1) were designed using Primer3 (https://bioinfo.ut.ee/primer3-0.4.0/), purchased from Integrated DNA Technologies (Coralville, IA), and confirmed by sequencing PCR products. Negative controls included reactions lacking reverse transcriptase or lacking template. Rps16 was used a reference gene for normalization and the delta-delta Ct method was used to calculate relative fold changes.

### RNA sequencing and bioinformatic analysis

Library preparation for RNA sequencing was performed by the Genomics Division of the Iowa Institute of Human Genetics. RNA samples were analyzed using an Agilent BioAnalyzer and samples with an RNA integrity number <8 were not used for sequencing. Total RNA libraries were prepared using the Illumina TruSeq Total RNA library preparation kit with rRNA depletion using the Illumina Ribo-Zero Gold Epidemiology kit. RNA sequencing was performed on an Illumina HiSeq 4000 platform to generate paired-end 150 bp reads. Each biological replicate was sequenced on three lanes, resulting in three technical replicates for each biological replicate.

Pre-processing, quality metrics, and alignment of sequence data was done on the University of Iowa Argon High Performance Computing cluster running CentOS-7.4 Linux. Quality metrics on the raw reads (in fastq format) were acquired using FastQC v0.11.8 (https://www.bioinformatics.babraham.ac.uk/projects/fastqc/). After the quality control assessment, technical replicates for each sample were merged to provide one fastq file for each read pair of each biological replicate. Trimmomatic v0.38 (http://www.usadellab.org/cms/?page=trimmomatic) was used to remove adapter sequences, using the adapter reference file provided by Trimmomatic, and trim all reads to a length of 100 base pairs to eliminate varying read length (`PE –threads 8 –phred 33 ILLUMINACLIP:TruSeq3-PE.fa:2:30:10 CROP:100`). Preliminary testing revealed that using a read length of 100bp provided optimal mapping percentage. After trimming, QC of the trimmed reads was assessed using FastQC to ensure adequate quality of trimmed reads prior to sequence alignment.

Sequence alignment was performed using STAR 2.7.0b (https://github.com/alexdobin/STAR, (Dobin et al., 2013)). Sequences were aligned to the rat genome (Rattus_norvegicus.Rnor_6.0.dna.toplevel.fa.gz) and annotated using Rattus_norvegicus.Rnor_6.0.95.gtf. Both the genome and annotation were downloaded from Ensembl. The genome index was generated using the following parameters:

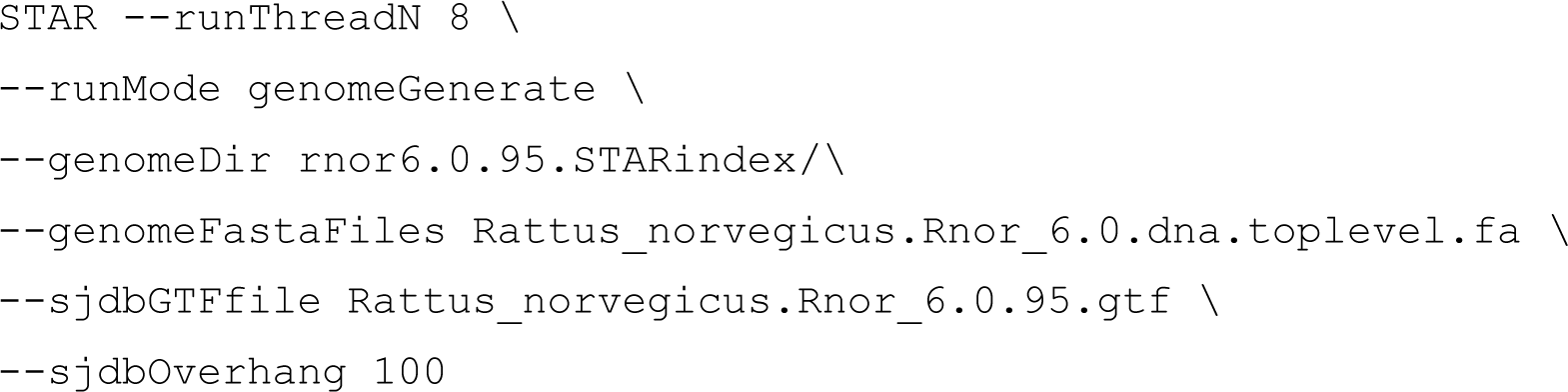

Reads were aligned using the following parameters:

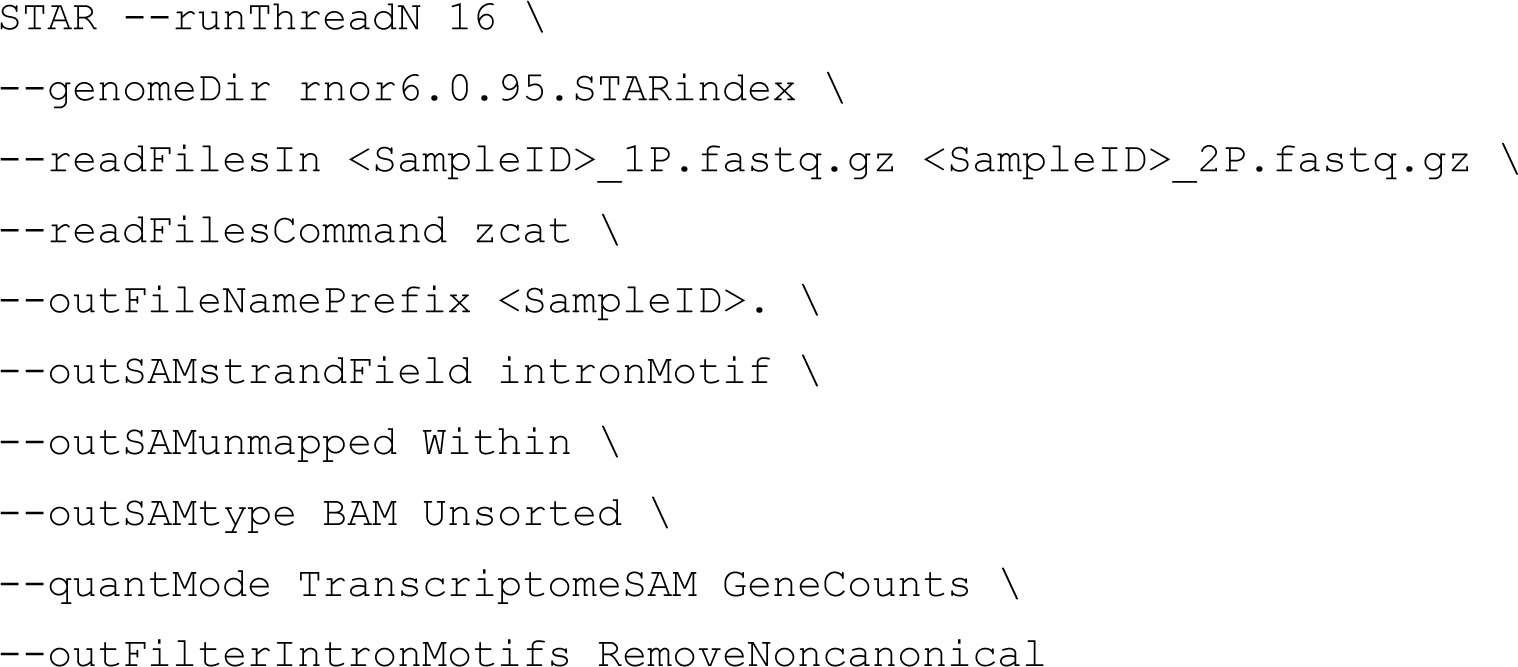

Mapped and unmapped reads were split into separate BAM files using samtools. Mapped reads were counted using *featureCounts* from the *subread* package (v1.6.3), using using Rattus_norvegicus.Rnor_6.0.95.gtf as a reference annotation file and the following parameters:

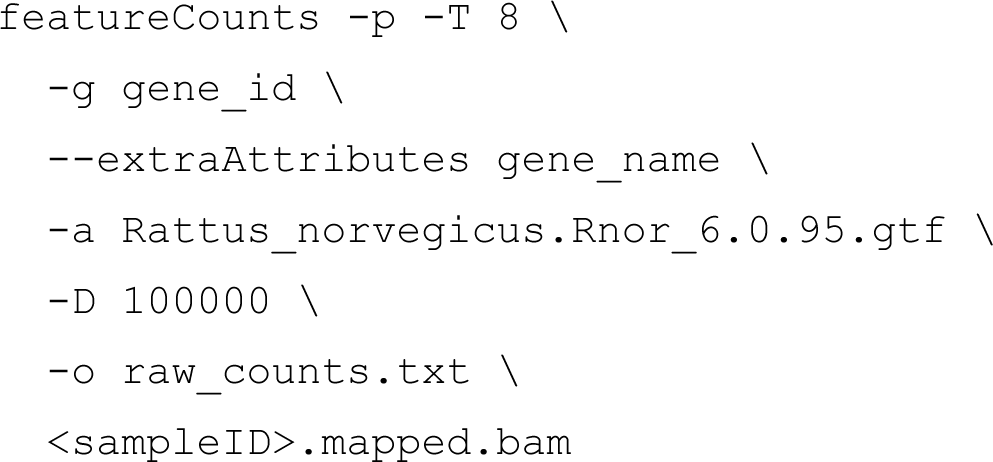

Raw counts were imported into R (v3.5.1 – v4.1.0) for further analysis. DESeq2 (1.32.0 (Love et al., 2014) was used to analyze differential expression of genes among the four conditions; P32 hearing (P32H), P32 deaf (P32D), P60 hearing (P60H), and P60 deaf (P60D). First, lowly expressed genes (defined as having an average raw count lower than ∼2-3 in all samples) were filtered out. Normalization of read counts and differential expression statistics were calculated using DESeq2. The cutoff for differential expression was an adjusted p-value (padj) < 0.05. Of these, genes with a greater than 2-fold change (log_2_FC) > 1 were considered upregulated and genes with a log_2_FC < -1 were considered downregulated. Results files of genes differentially expressed between hearing and deafened samples were generated to be used for downstream analysis. For heatmap visualization of gene expression of select gene sets, regularized logarithm normalization of counts was first calculated using the *rlog* function of DESeq2. Gene set matrices were then generated from the normalized count object and values were further normalized by row (gene) means. Heatmaps were generated using the ComplexHeatmap (v2.9.3) package in R. A principal component’s analysis (PCA) to ensure biological replicates clustered together, was done in R using DESeq2 functions and the package pcaExplorer (v2.18.0). The code for all analyses done in R is available upon request as a .rmd file.

### Gene set enrichment analysis on differentially expressed genes

Gene set enrichment analysis (GSEA) was done using GSEA software v4.1.0, (Mootha et al., 2003; Subramanian et al., 2005)) from http://software.broadinstitute.org/gsea/index.jsp to identify over-represented classes of genes in deafened versus hearing rats (and vice versa), overall and P32 and P60 separately. The most up-to-date gene ontology (GO) biological processes gene set (c5.bp.v7.0.symbols.gmt) from the Molecular Signatures Database was used. GSEA results were than used as input to Cytoscape (v3.8.2, (Shannon et al., 2003)) to visualize results (generation of networks and other plots). Networks were generated using the EnrichmentMap (v3.2.0, (Merico et al., 2010)) plugin. Only significant GO terms (nodes) (GSEA p<0.005 or False Discovery Rate Q<0.1) and significant edges (combined Jaccard and Overlap coefficient ≤ 0.375), which are a measure of overlap or similarity with other terms, were included in the networks. Based off the clusters formed by EnrichmentMap, the networks were manually edited/curated in Cytoscape to separate and modify clusters as well as define a broad category for each cluster using AutoAnnotate (DOI: 10.5281/zenodo.594859). The networks were then manually curated/edited for visual representation.

To perform GO analyses on specific sets of genes, the gprofiler2 package in R (Raudvere et al., 2019; Kolberg et al., 2020) was performed. gProfiler, in combination with Cytoscape and EnrichmentMap, was used for classifying functional groups of genes, as described in Results. Manhattan plots of gProfiler results were generated in R using the ggplot2 package.

### Determination of auditory thresholds using auditory brainstem response (ABR) measurements

ABR measures were performed at or close to P32 in all rats. Responses were recorded in from 90 dB SPL to 10 dB below threshold in 10 dB steps, repeated in 5 dB steps near threshold. The lowest stimulus level at which there was a detectable wave I was considered threshold. ABRs were recorded using an RZ6 multi I/O processor with a RA4PA 4 channel preamplifier (Tucker-Davis Technologies, Inc.) in a custom-made sound-proof chamber. BioSigRZ (version 5.6, Tucker-Davis Technologies, Inc.) was used to generate and deliver the stimuli and record response signals. The acquisition period was 12 ms with a sampling rate of 25,000/s. A high-pass filter was set at 3000 Hz and the low-pass filter at 300 Hz. All final signals are an average of 512–1024 sweeps.

Rats were anesthetized with a mixture of ketamine (40 mg/kg) and xylazine (4 mg/kg). Our criterion for success in eliminating hair cells was histological observation of hair cell loss in one cochlea from each rat. Nevertheless, we assessed auditory brainstem response (ABR) to exclude from this study, at an early point, rats that not completely deafened. We measured ABR in all rats at P32 (click stimuli, 1-70 kHz, 90 to 40 dB SPL in 10 dB increments). Any kanamycin-treated rats with detectable ABR wave 1 for stimuli of 90 decibels sound pressure level (dB SPL) was excluded from further study.

To assess hearing in C3KO rats, responses to tone pips with a duration of 5 ms and gated time of 0.5 ms, presented at a rate of 21/s, alternating polarities at 4, 8, and 16 kHz were measured in normal hearing wildtype, C3 heterozygous, and C3KO rats. Tone pip stimuli were delivered to the external auditory meatus via a custom-made insertion tube connected to an MF1 speaker (Tucker-Davis Technologies, Inc.) by a polyethylene tube. Responses were measured from 90 dB SPL to 10 dB below threshold in 10 dB steps, and repeated in 5 dB steps near threshold. The lowest stimulus level at which there was a detectable wave I was considered threshold. The recording electrode configuration for all rats consisted of an active electrode placed at the midline of the vertex of the skull, a reference electrode at the ipsilateral mastoid area, and a ground electrode in the low blank/flank area.

ABR data were exported from BioSigRZ and off-line processing and analysis was performed using custom-written scripts in MATLAB (R2021a). Wave I amplitude was determined by measuring the difference (in µV) between the wave I peak and the following negative trough. Wave I amplitude growth curves as a function of stimulus intensity were fit by a second-order polynomial function for group comparisons. Wave I latency was defined as the time, in ms, from the start of the stimulus (0 ms) to the wave I peak. The MATLAB scripts used for processing and analysis are available at https://github.com/bgansemer/ABR-analysis.

### Immunohistochemistry

Tissue collection and processing for fluorescent labeling of neurons and immune cells was done as previously described (Rahman et al., 2023). Rats were anesthetized with ketamine / xylazine, as above, and, while under surgical plane, transcardially perfused with ice-cold PBS followed by ice-cold 4% paraformaldehyde (PFA) in 0.1 M phosphate buffer. Cochleae were isolated from the temporal bones and post-fixed in 4% PFA at 4°C for 16-24 h, followed by decalcification for 5-10 days in 0.12 M EDTA with daily changes of EDTA. The cochleae were then cryo-protected by 30 min incubations each of 10, 15, 20, 25, and 30% sucrose in PBS, then rotated in degassed O.C.T. compound (Tissue-Tek) overnight at 4°C. The cochleae were infiltrated and degassed in O.C.T. under gentle vacuum for 1-2 h, embedded in O.C.T., and flash-frozen in liquid nitrogen. Embedded cochleae were stored at −80°C until sectioning. Serial sections were cut parallel to the midmodiolar plane on a Leica Cryostat CM1850 and collected on Fisher Superfrost slides. Sections were cut at 25 µm thick for and stored at −80°C until immunolabeling was performed.

Prior to immunolabeling, slides were warmed to room temperature (∼20-22°C). Cochlear sections were then permeabilized with 0.5% Triton X-100 in Tris-buffered saline pH 7.6 (TBS) for 25 min, washed with TBS, and immersed in blocking buffer (5% goat serum (company), 2% bovine serum albumin (Sigma, catalog #A2153), 0.1% Triton X-100, 0.3% Tween-20, and 0.02% sodium azide (Sigma, catalog #S2002) in TBS) for 3-4 h at room temperature. After blocking, sections were incubated in primary antibody in blocking buffer overnight (∼16 h) at 4°C. After primary antibody application, sections were washed in TBS then incubated in blocking buffer containing secondary antibodies (Alexa Fluor conjugates, 1:400, Invitrogen) for 3-4 h at room temperature. Sections were then washed in TBS and nuclei stained with Hoechst 3342 (10 µg/ml in PBS, Sigma). Slides were then washed and coverslipped with Fluoro-Gel Mounting Medium with Tris Buffer (catalog #17985-10, Electron Microscopy Sciences).

For cochlear wholemounts, the cochleae were prepared as above until the sucrose gradient step. After decalcification, microdissection of the cochleae was performed to remove extraneous tissue and ensure the decalcified bony capsule was sufficiently open to allow infiltration of labeling solutions. The whole cochleae were permeabilized in 1% Triton X-100 in PBS for 1 h at RT, followed by 3x5 min washes in PBS. Cochleae were then immersed in blocking buffer (5% goat serum, 2% BSA, 0.1% TritonX-100, 0.02% sodium azide) for 1-2 h at RT. After blocking, cochleae were incubated in primary antibody in blocking buffer for 48-72 h at 4°C. The cochleae were then washed 3x with PBS, post-fixed for 2 h in 4% PFA, washed 3x with PBS, and then incubated in secondary antibody in blocking buffer for 24 h at 4°C. After the secondary antibody incubation, the cochleae were dissected for whole mount preparation. The basilar membrane with the organ of Corti and associated spiral ganglion were cut into 5-6 pieces and mounted on a glass slide and coverslipped with Fluoro-Gel Mounting Medium.

Primary antibodies used in this study included mouse IgG2a-anti-β-III-tubulin (Tuj1, 1:400, #801202, Biolegend) to label SGNs, rabbit-anti-Iba1 (1:800, #019-19741, Wako) to label macrophages, mouse IgG1-anti-CD68 (ED1, 1:400, #31360, Abcam) to label activated macrophages, combined rabbit-anti-myosin VI (1:400, #KA-15, Sigma) and rabbit-anti-myosin VIIa (1:400, #25-6790, Proteus) to label hair cells, mouse IgG1-anti-rat CD45 (OX-1, 1:100, #202201, Biolegend) to label CD45+ cells, and mouse IgG1-anti-rat MHCII (OX-6, 1:100, # 50-165-477 Biolegend) to label MHCII+ cells.

### Imaging and quantitation

Imaging and quantitation were performed as previously described for 25 µm thick sections (Rahman et al., 2023). Labeled sections were imaged on a Leica SPE confocal system using a 40X (0.95 NA) objective, 1x digital zoom, and a z-axis spacing of 1 µm. Higher magnification images were taken using a 63X (1.3 NA) objective, 2-2.5x digital zoom, and z-axis spacing of 0.5 µm. Gain and contrast settings remained constant throughout imaging sessions to maintain consistency. Fiji/ImageJ (NIH, https://imagej.net/Fiji) was used for all image analysis and all cell counts were done on maximum intensity z-projections of the image stacks. Using an automated code generator written in Python, all images were assigned an 8-digit code number to ensure that all analyses were done blind to the experimental condition from which images came.

For all counting, the outline of Rosenthal’s canal was traced and the cross-sectional area measured in order to calculate cell density per mm^2^ for all cell counts. For immune cells, macrophages, CD45+ lymphocytes, and MHCII+ cells were counted using anti-Iba1, anti-CD45, and anti-MHCII antibodies, respectively. All Iba1+ cells within Rosenthal’s canal were counted to determine macrophage density. The number of MHCII+ macrophages (Iba1+/MHCII+) was also determined. For CD45+ and MCHII+ cells, sections were labeled with anti-Iba1 and either anti-CD45 or anti-MHCII. Because macrophages (Iba1+ cells) were found to be immunoreactive for CD45, the Iba1+ signal was first subtracted from the CD45 signal to ensure that double positive (Iba1+/CD45+) were not counted and only Iba1-/CD45+ cells were counted. To determine SGN survival, sections were labeled with anti-β-III-tubulin (TuJ1). Because SGN diameter is less than 25 µm and multiple layers of SGN soma could be present in 25 µm thick sections, “virtual 6 µm sections” were created by taking the middle 7 z-slices from each z-stack. All SGN counts were performed from these virtual thin sections. For all counts, only cells with a visible nucleus were counted.

For all histological analyses, cochlear location was defined by half-turn increments proceeding from the base to the apex as previously described (Kopelovich et al., 2013). Cell densities from each cochlear location for an animal were averaged together to provide one “n”. Thus, “n” is the number of rats used in the study. Control and experimental conditions were imaged in the same sessions using identical settings to allow quantitative comparison.

### Western blotting

At the time of tissue collection for histology, prior to perfusion with paraformaldehyde, a small portion of liver tissue was collected in a microfuge tube and frozen on dry ice. Samples were stored at −80°C until processing. The liver tissue was thawed and lysed in RIPA buffer (150 mM NaCl, 5 mM EDTA, 50 mM Tris, 1% NP-40, 0.5% sodium deoxycholate, and 0.1% SDS) and homogenized with a pellet pestle (RPI, #199228). Protein lysates were diluted with Tris-SDS buffer (100 mM Tris, 1% SDS) and boiled for 15 min. Protein concentration was measured using a Pierce BSA Protein Assay Kit following manufacturer instructions (Pierce, #23277). Equal amounts of protein lysate (∼10 µg per lane) from each sample were separated on 10% Tris-HCL SDS-PAGE gels and transferred to a nitrocellulose membrane (0.45 µm pore size, #LC2001, Invitrogen). Blots were blocked with fish gelatin (Biotium, #22010) in TBS for 1 h at RT and then probed with anti-C3 (1:2000, #0855730, MP Biomedical) in fish gelatin in TBS + 0.1% Tween-20 (TBST) for 2 h at RT. Primary antibody against GAPDH (1:5000, #CB1001, Millipore) was used as a loading control. Blots were then washed with TBST and incubated with near infrared dye-conjugated secondary antibodies (1:20,000, Invitrogen) in fish gelatin in TBST). Blots were visualized on a Licor Odyssey FC imaging system.

### Statistical analysis

R (v3.5.1 – v4.1.0) was used for statistical analysis of the RNAseq data as described above. Graphpad Prism (version 9.2.0) was used for statistical analysis of all cell count data and to generate graphs of cell count data. The ABR data collected from MATLAB were further analyzed in Prism to perform curve fitting and generate graphs of averaged data. For comparisons with a multiple groups, a one-or two-way ANOVA with Tukey’s multiple comparison was used for parametric data and a Kruskal-Wallis with Dunn’s multiple comparisons was used for nonparametric data, as indicated in figure legends. For pairwise comparisons, a Mann-Whitney U test was performed. Multiple unpaired t-tests with Benjamini, Kreiger, and Yekutieli false discovery rate for multiple comparisons was used for statistical analysis of qPCR data. A p<0.05 was considered statistically significant.

### Data availability

The RNAseq gene expression data are available in an easily viewable and interactive format on gEAR (www.umgear.org, “RNA-seq analysis of the spiral ganglion of hearing and aminoglycoside-deafened rats”). All raw RNAseq data, along with spreadsheets of normalized data, are available at GSE194063. Analysis files and scripts/code are available from the authors upon reasonable request.

## Results

### RNAseq profiling of the spiral ganglion of hearing and deafened rats

We used bulk RNAseq to assess gene expression in the spiral ganglia of hearing and deafened rats at P32 and P60. As in our previous study (Rahman et al., 2023), these time points were chosen because P32 is approximately two weeks after the last inner hair cells die (Bailey and Green, 2014) but prior to onset of most SGN death, while by P60 nearly half of the SGNs have died.

In assessing the quality of the RNAseq data, an initial principal component analysis using the 750 most variant genes and plotting principal component (PC) 1 and PC2 revealed that the third biological replicate of the P32H condition did not cluster with the other two biological replicates (Supplementary Figure 2). Therefore, this biological replicate was removed and the differential expression analysis re-run without the replicate. All subsequent analyses were done using the two remaining P32H replicates and all three biological replicates from the other conditions.

After filtering out low-read-count genes such that any gene with an average of <1 read per sample was excluded, there were 21,829 genes found to be expressed in the spiral ganglion. Of these 21,829 genes, 4,346 were significantly differentially expressed defined as a DESeq2 adjusted p value (padj) <0.05 between deafened and hearing samples. This includes genes differentially expressed at P60, at P32 or at both times. Of these, 598 genes had a log_2_-fold change (log_2_FC) >1 (upregulated) and 251 had a log_2_FC<-1 (downregulated). (This corresponds to an increase >2-fold or a decrease to <0.5.) Regarding normal maturational changes, 164 genes were significantly differentially expressed between P60H and P32H, with 12 being upregulated and 49 downregulated. Table 1 summarizes the number of genes that are significantly differentially expressed and upregulated or downregulated for each individual comparison. As shown in Figure 1B, when comparing gene expression between hearing and deaf there are both up-and down-regulated genes in the top 500 most significant genes (lowest padj values). Of the top 500 most significant genes, 251 had increased (log_2_FC>0) and 249 had decreased (log_2_FC<0) expression after deafening.

**Table 1.**
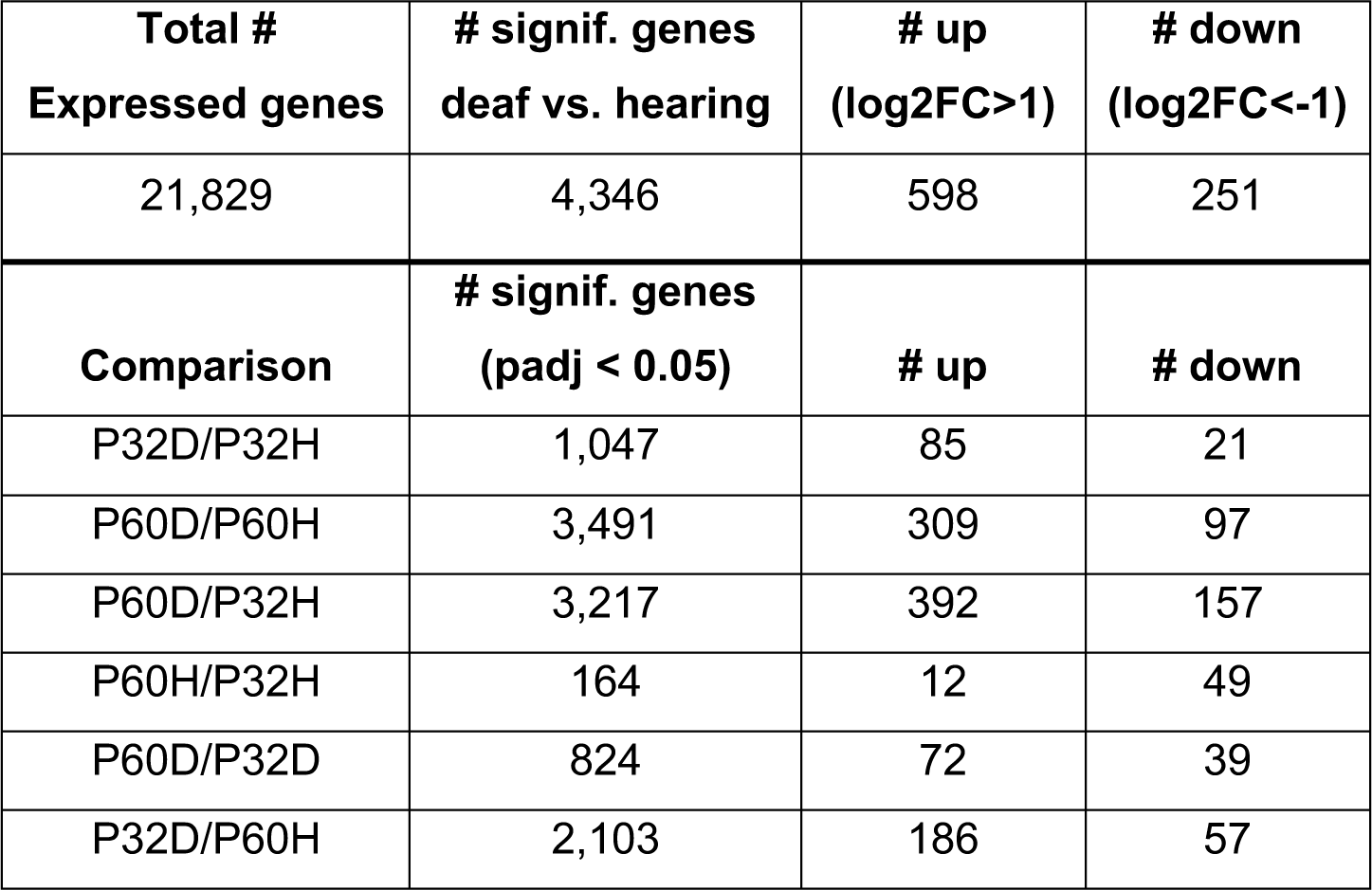
Number of significantly differentially expressed genes by comparison. The number of expressed genes with number of genes with a padj <0.05 for each comparison was determined using DESeq2. The Columns 2 and 3 show the number of significant genes for each comparison that are upregulated (log2FC > 1) or downregulated (log2FC < -1). padj: adjusted p-value; log2FC: log2 fold change; P32D: P32 deaf; P32H: P32 hearing; P60D: P60 deaf; P60H: P60 hearing

| **Total #**<br>**Expressed genes** | **# signif. genes**<br>**deaf vs. hearing** | **# up**<br>**(log2FC>1)** | **# down**<br>**(log2FC<-1)** |
| --- | --- | --- | --- |
| 21,829 | 4,346 | 598 | 251 |
| **Comparison** | **# signif. genes**<br>**(padj < 0.05)** | **# up** | **# down** |
| P32D/P32H | 1,047 | 85 | 21 |
| P60D/P60H | 3,491 | 309 | 97 |
| P60D/P32H | 3,217 | 392 | 157 |
| P60H/P32H | 164 | 12 | 49 |
| P60D/P32D | 824 | 72 | 39 |
| P32D/P60H | 2,103 | 186 | 57 |

We suggest that there can be two possible reasons for expression of a particular gene to differ between hearing and deafened animals at P60. One is that a gene is normally stably expressed after P32 but, because of hair cell loss, is up or down regulated. The second applies to genes that are normally up-or down-regulated during maturation (and there are many such genes, as will be described below). For some of these, deafening prevents the normal maturational change and expression of the gene at P60 will remain similar to that at P32 instead of being decreased or increased as normal. For example, a gene that normally increases in expression level between P32 and P60 (e.g., Syt7) but fails to do so in deafened rats would have significantly lower expression in P60D ganglia relative to P60H ganglia and would be initially classified as “downregulated due to deafening” at P60. Conversely, a gene that normally decreases in expression level between P32 and P60 (e.g., Fabp5) but fails to do so in deafened rats would have significantly higher expression in P60D ganglia relative to P60H ganglia and would be initially classified as “upregulated due to deafening” at P60. To distinguish whether an apparent up-or down-regulation at P60 is a direct response to deafening or due to the deafening preventing normal maturational change, we compared the ratio of P60D expression to P32H expression (P60D/P32H). If this ratio is close to 1 (-0.5 < P60D/P32H log_2_FC < 0.5), we infer that deafening prevented up or down regulation and classify any difference in expression between P60H and P60D accordingly.

Based on a criterion of padj<0.05, expression of 3,491 genes was found to be significantly different between P60D and P60H. Of these, expression of approximately half, 1,769 genes, was significantly different, but with only a modest change in expression level (-0.5 < P60D/P60H log_2_FC < 0.5). Of the remaining 1,722 genes, 913 were classified as upregulated post-deafening (P60D/P60H log_2_FC>0.5) and 809 downregulated post-deafening (P60D/P60H log_2_FC<-0.5). These were further subdivided based on the value of P60D/P32H log_2_FC: 750 genes for which upregulation was due directly to deafening, 163 genes for which normal downregulation between P32 and P60 was prevented by deafening, 602 genes for which downregulation was due directly to deafening, 207 genes for which normal upregulation between P32 and P60 was prevented by deafening. Note that we use a less stringent criterion for up-or downregulation than we used for Table 1. This was to increase the number of genes in the sets and so allow a more accurate comparison and distinction among the different gene sets. These gene sets were then used as input to gProfiler for gene ontology analyses.

### Gene set enrichment analysis of RNAseq gene expression data

To identify possible functionally related groups of genes that were either up-or down-regulated after deafening, the most significantly (lowest padj) upregulated (251 genes) or downregulated (249 genes) were used as input to gProfiler package in R using the `gost` function. Figure 1C shows Manhattan plots of the results from the downregulated (top) or upregulated (bottom) genes. In each plot, representative categories with the highest -log10 (p value) are highlighted. These data will be described in later sections. To perform a more unbiased GO analysis, GSEA was performed on the complete set of normalized read counts to compare deafening-related changes overall and at P32 and P60 separately. Figure 1D is a network visualization of the overall deafened vs. hearing changes. A corresponding graph depicting the number of gene ontology categories in each cluster is shown in Supplemental Figure 3A. Overall, 647 gene ontology categories (nodes) were identified as enriched: 557 in deafened (red nodes) and 90 in hearing (blue nodes). Clusters of categories were determined by similarity between categories; categories that shared genes were clustered together. This resulted in formation of 15 clusters, 12 enriched in deafened and 3 in hearing. Of note, two of the clusters enriched in hearing, synaptic transmission (26 nodes) and ion transport/homeostasis (40 nodes), contain gene ontology categories mostly associated with neuronal function. Several clusters were identified for categories enriched in deafening. These include apoptosis (19 nodes), phagocytosis (9 nodes), cell differentiation/migration (38 nodes), and cell adhesion (21 nodes), among others. Most prominently, the largest cluster contained categories associated with the immune response (292 total nodes, 52.4% of nodes enriched in deafened). Moreover, this large immune response cluster could be further subdivided in smaller clusters as shown in Figure 1D and Supplemental Figure 3A. This indicates that most upregulated genes are related to an immune/inflammatory response.

Network visualizations of deafening-related changes at P32 and P60 separately are shown in Figure 2. The corresponding graphs showing the number of categories in each cluster are shown in Supplemental Figure 3, with immune response cluster names also identified in Table 2. Many of the same clusters/categories were identified as in the overall analysis, and there were slightly more categories identified as enriched when comparing deafened to hearing at P60 (547 nodes) than at P32 (517 nodes). One cluster that was not present at P32 but only at P60 was the apoptosis category. This is expected as neuronal apoptosis is more active at P60 than at P32 (Alam et al., 2007). At both timepoints, the categories identified as enriched in hearing were largely associated with neuronal/synaptic structure and function, suggesting that changes intrinsic to SGNs are occurring after deafferentation, and could be occurring as early as P32. Finally, like the overall hearing vs. deafened analysis, most categories enriched in deafened were associated with immune and inflammatory responses. This result implies that, while the period of SGN death is only just beginning at P32, multiple immune response pathways have already been activated, and continue to be activated at P60. This is consistent with the immune response being causal to SGN death rather than a response to SGN death.

**Figure 2.**
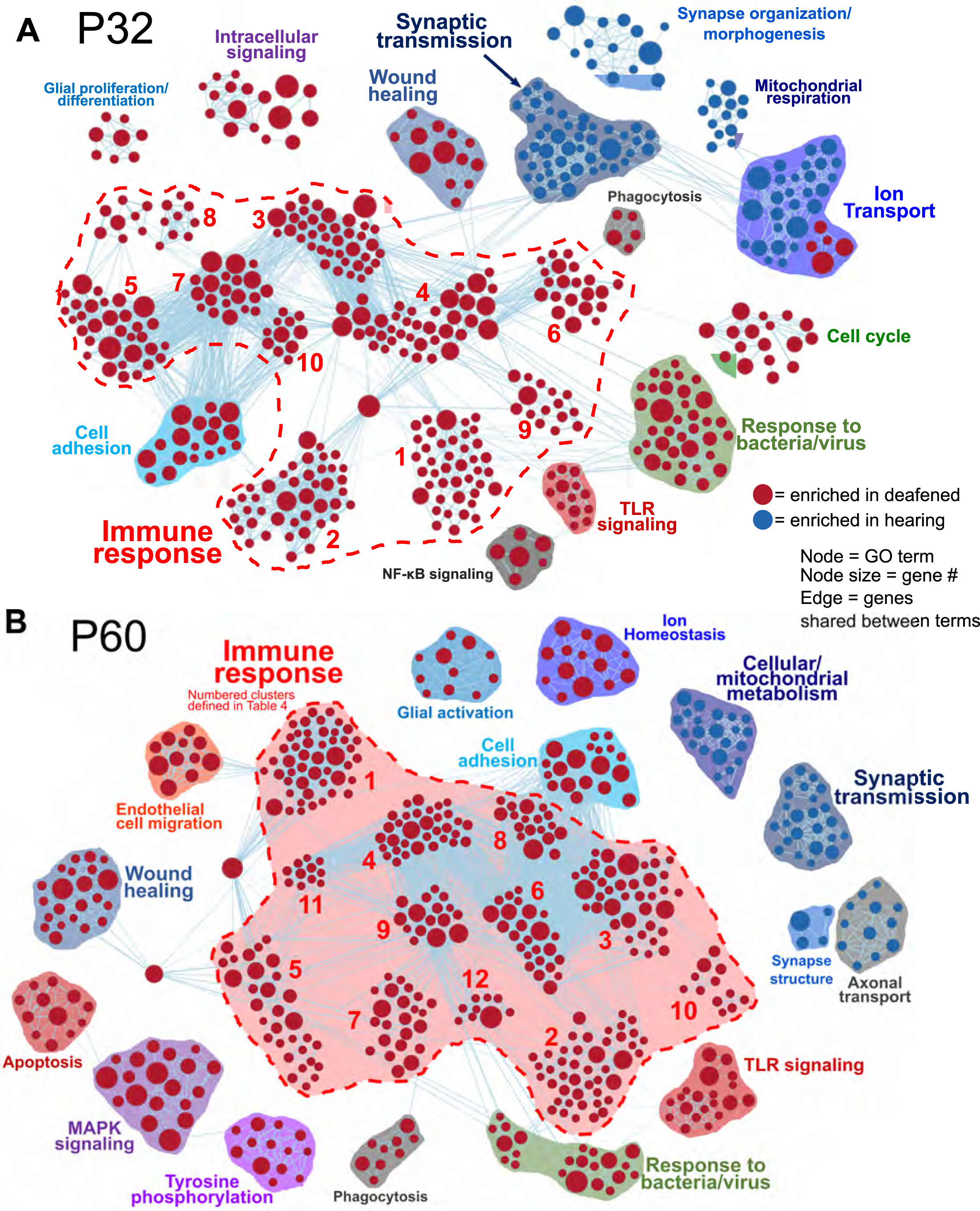
Comparative analysis of hearing vs. deafened separately at P32 and P60. GSEA networks showing enriched categories when comparing hearing vs. deafened at P32 alone (A) or P60 alone (B). Red nodes are GO categories enriched in deafened, blue nodes are those enriched in hearing. Edges (lines) indicate shared genes between the two connected terms. The category names for the numbered immune response clusters are defined in Table 2 – and are the same as used in Supplemental Figures 3B and 3C.

**Table 2.**
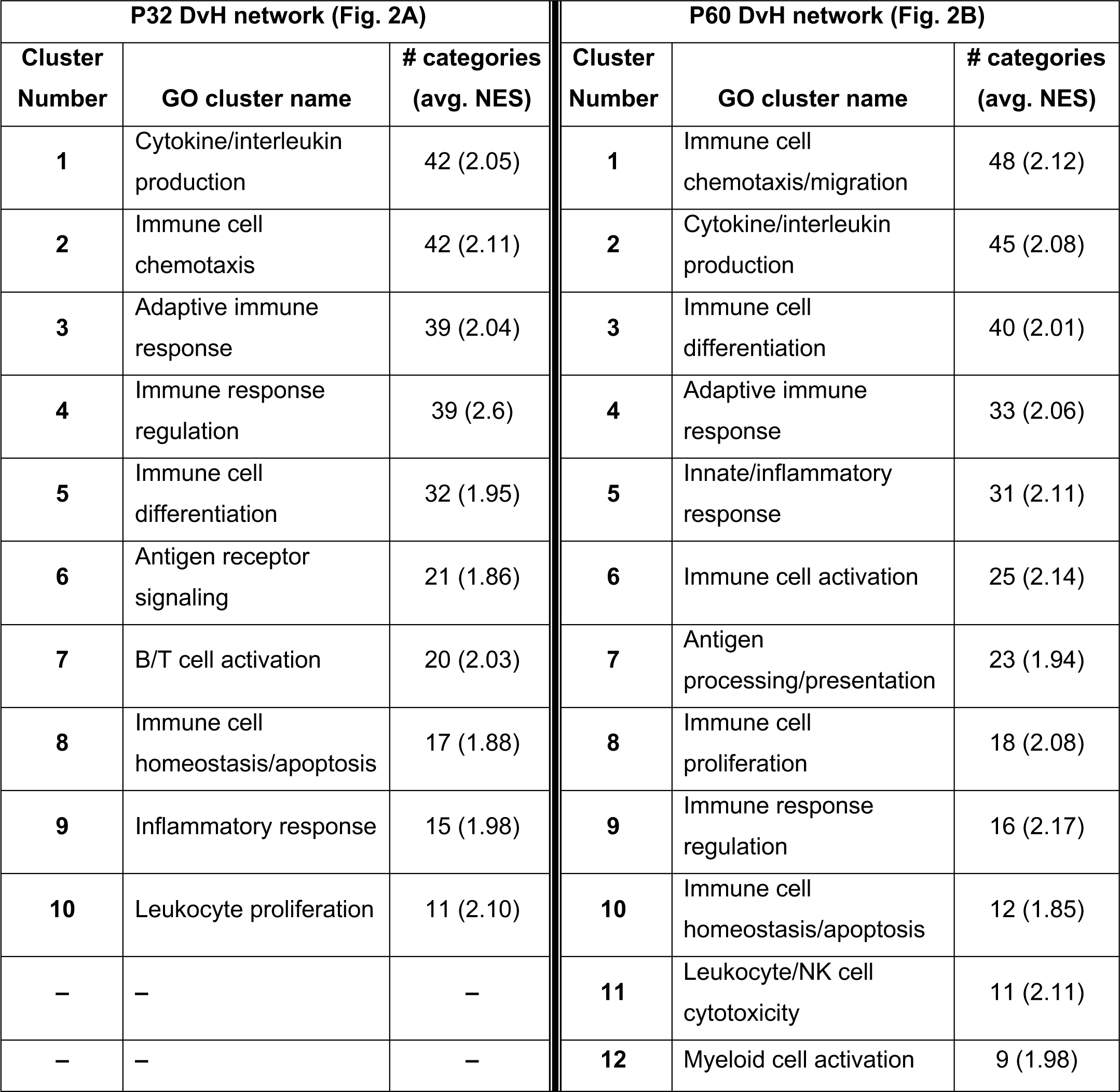
Key to name and normalized enrichment score (NES) of immune response gene clusters indicated by numbers in figures 2A and 2B. Higher NES indicates greater activation of the genes in the cluster.

| P32 DvH network (Fig. 2A) |  |  | P60 DvH network (Fig. 2B) |  |  |
| --- | --- | --- | --- | --- | --- |
| Cluster Number | GO cluster name | # categories (avg. NES) | Cluster Number | GO cluster name | # categories (avg. NES) |
| 1 | Cytokine/interleukin production | 42 (2.05) | 1 | Immune cell chemotaxis/migration | 48 (2.12) |
| 2 | Immune cell chemotaxis | 42 (2.11) | 2 | Cytokine/interleukin production | 45 (2.08) |
| 3 | Adaptive immune response | 39 (2.04) | 3 | Immune cell differentiation | 40 (2.01) |
| 4 | Immune response regulation | 39 (2.6) | 4 | Adaptive immune response | 33 (2.06) |
| 5 | Immune cell differentiation | 32 (1.95) | 5 | Innate/inflammatory response | 31 (2.11) |
| 6 | Antigen receptor signaling | 21 (1.86) | 6 | Immune cell activation | 25 (2.14) |
| 7 | B/T cell activation | 20 (2.03) | 7 | Antigen processing/presentation | 23 (1.94) |
| 8 | Immune cell homeostasis/apoptosis | 17 (1.88) | 8 | Immune cell proliferation | 18 (2.08) |
| 9 | Inflammatory response | 15 (1.98) | 9 | Immune response regulation | 16 (2.17) |
| 10 | Leukocyte proliferation | 11 (2.10) | 10 | Immune cell homeostasis/apoptosis | 12 (1.85) |
| – | – | – | 11 | Leukocyte/NK cell cytotoxicity | 11 (2.11) |
| – | – | – | 12 | Myeloid cell activation | 9 (1.98) |

As previously mentioned, changes in gene expression between P60D and P60H ganglia could be due to a lack of a maturational process or new changes as result of deafening. Figure 3 shows network visualizations of gene ontology analyses for each of four gene sets: genes upregulated due to deafening (Figure 3A), genes apparently upregulated due to a lack of maturational downregulation (Figure 3B), genes downregulated due to deafening (Figure 3C), and genes apparently downregulated due to a lack of maturational upregulation (Figure 3D). There were 750 genes identified as upregulated as a new consequence of deafening. When used as input to gProfiler, 250 gene ontology categories were identified as enriched. These were organized into 15 clusters, the majority of which contained immune response-related categories. Prominent clusters include adaptive immune response (53 categories), cytokine/chemokine production (45 categories), antigen processing/presentation (14 categories), and leukocyte/lymphocyte activation (15 categories). This implicates adaptive immune response processes as potential contributors to SGN death post-deafening. The 163 genes that did not undergo maturational downregulation were in only 10 categories, all related to cell cycle and cell division. This, along with categories associated with cell proliferation (particularly immune cell proliferation) being identified in the deafened vs. hearing analyses (Figure 1D, 2, Supplemental Figure 3), suggests that decreases in cell division and proliferation that might normally occur from P32 to P60 do not occur after deafening, and cell division may even increase. This is consistent with a previous observation of increased Schwann cell division after deafening (Provenzano et al., 2011) but may also include proliferation of macrophages. When assessing genes downregulated in P60D vs. P60H, 602 were categorized as downregulated due directly to deafening. Gene ontology analysis identified 118 categories that grouped into 8 clusters. These categories were associated with neuronal and synaptic function. Similar, but smaller, clusters were identified when the 207 genes that failed to undergo maturational upregulation were used as input to gProfiler. This indicates that the majority of downregulatory changes are occurring in the neurons and are associated with neuronal functions and signaling. While these changes are not the major focus of this study, they will be briefly described in greater depth in the following section.

**Figure 3.**
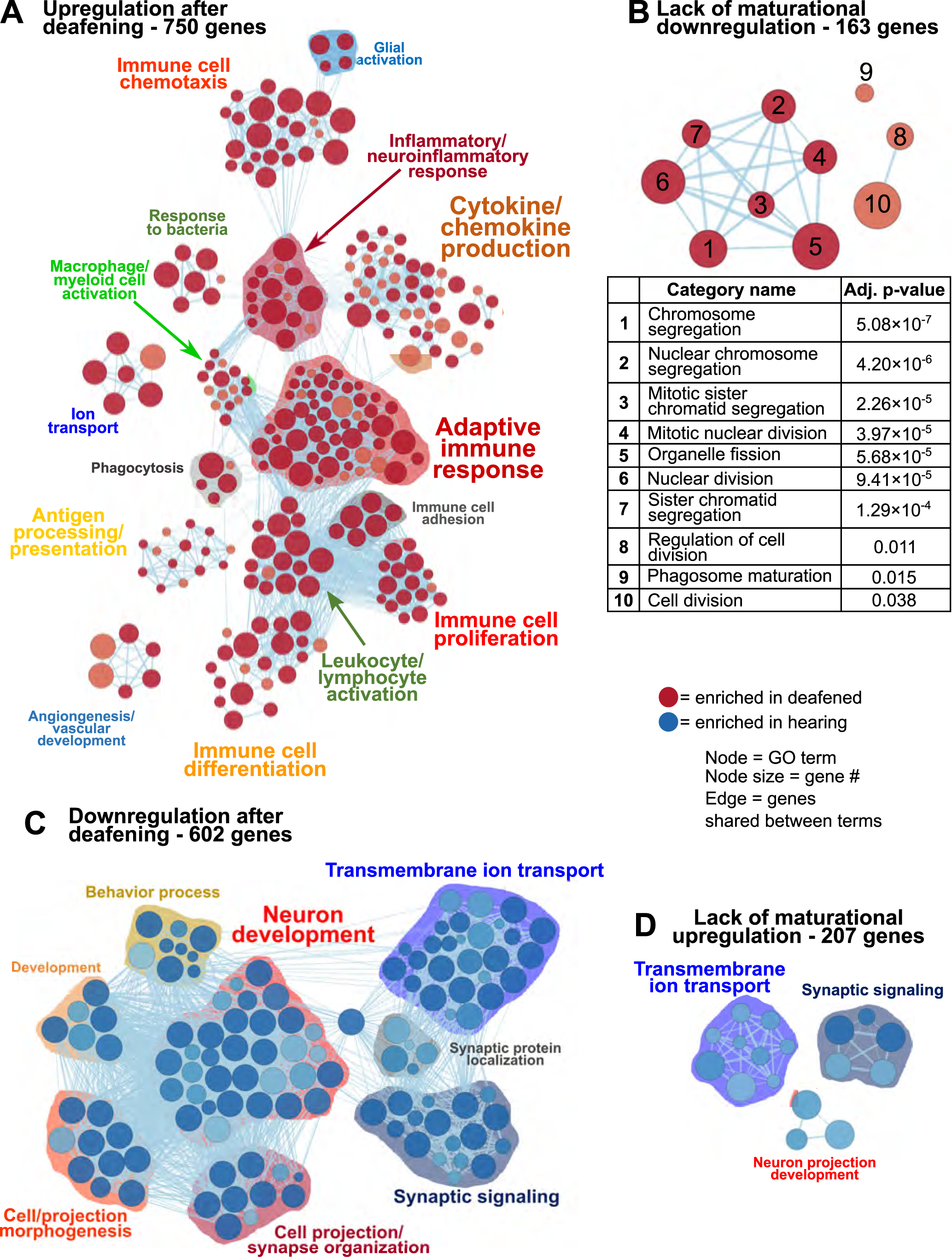
Differences in gene expression between deafened and hearing P60 rats. due directly to deafening or, indirectly, due to lack of the normal maturational change. As described in Results, genes upregulated or downregulated at P60D vs. P60H were classified as changed either due to deafening or due to a lack of a maturational change, based on the P60D/P32H ratio. These gene sets were used as input to gProfiler and network visualizations generated. Network visualizations showing GO terms enriched for genes upregulated due to deafening (A), upregulated due to a disrupted maturational downregulation (B), downregulated due to deafening (C), or downregulated due to a disrupted maturational upregulation (D) are shown. The table under B indicates the term names and adjusted p values for the terms in the network in B. Each node is a GO term and edges (lines) indicated shared genes between terms.

### Neuronal/synaptic genes are downregulated after deafening

The prominent clusters enriched in hearing in the GSEA and gProfiler analyses contained categories associated with synaptic transmission, axonal conduction, and other neuronal functions (Figure 1CD, 2, 3C). This suggests that the most significant transcriptional *down*regulation occuring after deafening is for genes that play roles in neuronal function. When using the 500 genes with greatest statistical significance to downregulation of their expression post-deafening as input to gProfiler, ten of fifteen categories with significant enrichment were associated with synaptic signaling or transmembrane transport (Figure 1C, Table 3). Additionally, while the GSEA identified some categories related to synaptic structure as downregulated post-deafening at both P32 and P60 (Figure 2), there were fewer categories than those pertaining to synaptic transmission/function (14 vs. 46 at P32 and 4 vs. 24 at P60). Of note, many of the downregulated genes were associated with presynaptic functions such as vesicle trafficking, vesicle tethering/fusion, and neurotransmitter release. These include several synaptotagmins (i.e., Syt2, Syt3, and Syt7), Cplx1, Rab3a, Rims1, Rims2, and Synj1 are all significantly downregulated after deafening at P32 and/or P60 (Figure 4D). Other categories with genes that were identified as significantly differentially expressed after deafening include voltage-gated ion channels (Figure 4C), Ca^2+^ pumps and binding proteins (Figure 4E), and neuronal transcription factors and cytoskeletal proteins (Figure 4B). Notably, channels involved in high frequency firing, a distinctive feature of auditory neurons, e.g., Kcnc1 and Kcnc3 are downregulated after deafening. NTFs expressed in the spiral ganglion continue to be expressed after deafening with either no change in level (e.g., Ntf3, Artn, and Nrtn), or increase in level (Bdnf and Cntf). Also, cognate NTF receptors expressed in the spiral ganglion – Ntrk2 (TrkB), Ntrk3 (TrkC), Ret, Gfra2, and Cntfr – show no significant change in expression after deafening. Overall, these results suggest that, after deafening, the ability of SGNs to carry out their characteristic neuronal functions, particularly at their central synapses, is declining due to downregulation of key genes, although this is not because of loss of neurotrophic support.

**Figure 4.**
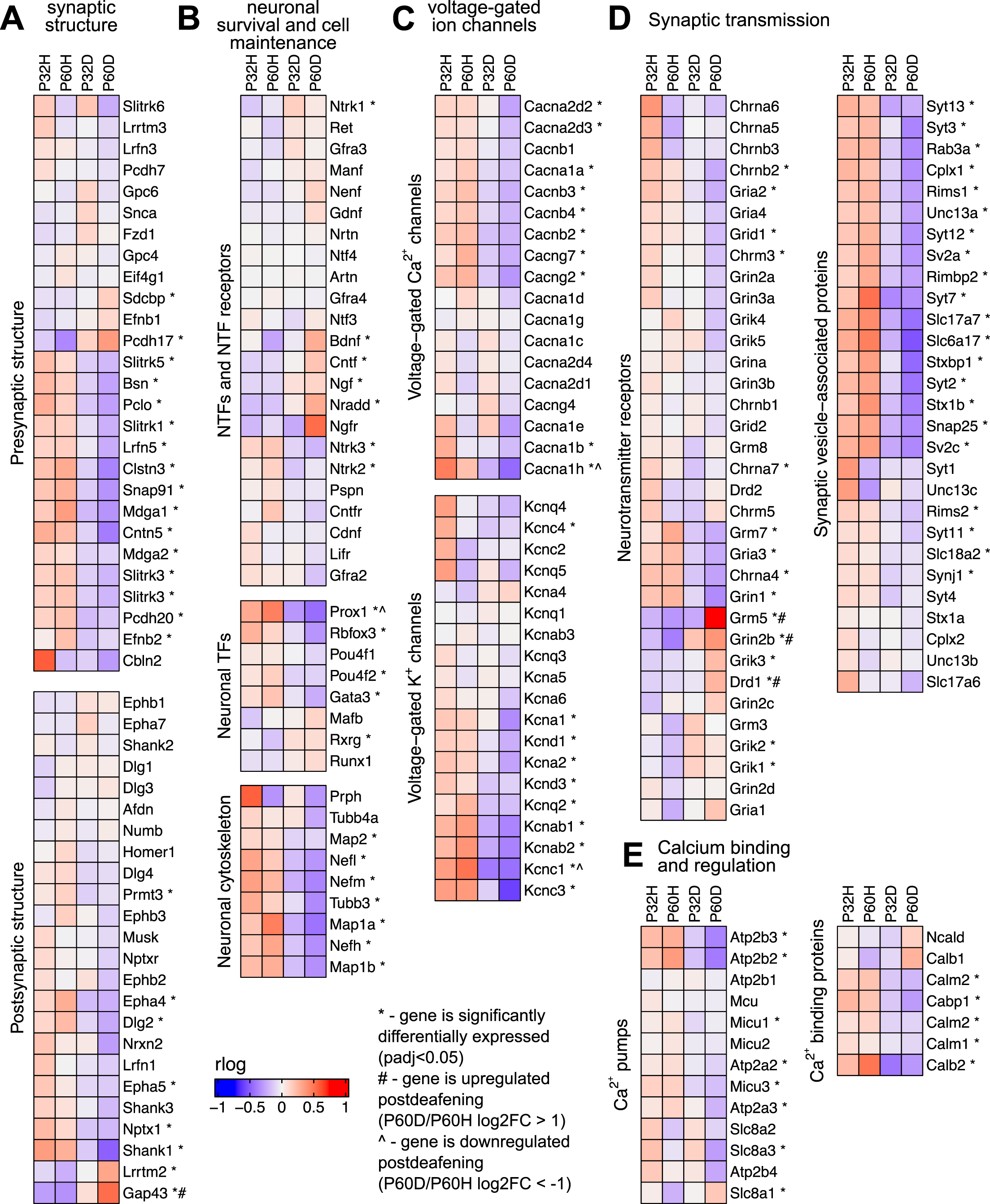
Neuronal/synaptic genes are downregulated after deafening. A) Heatmap showing regularized logarithm (rlog) normalized expression of genes involved in presynaptic (top) or postsynaptic (bottom) structure across conditions. B) Heatmap of genes for NTFs and NTF receptors (top), neuronal transcription factors (middle), and neuronal cytoskeletal proteins (bottom). C) Heatmap of genes for voltage-gated Ca^2+^ channels (top) and voltage-gated K^+^ channels (bottom). D) Heatmap of genes for neurotransmitter receptors (left) or genes for synaptic vesicle-associated proteins (right). E) Heatmap of genes for Ca^2+^ pumps (left) and Ca^2+^ binding proteins (right). The expression shown for each condition is the averaged normalized expression of the biological replicates for that condition.

**Table 3.** Genes with lower expression at P60 in deafened relative to hearing (P60D/P60H) GO terms associated with axonal or synaptic transmission are highlighted.

| GO term ID | GO term name | p value | term size | intersection |
| --- | --- | --- | --- | --- |
| GO:0007268 | chemical synaptic transmission | 2.59E-05 | 264 | 15 |
| GO:0098916 | anterograde trans-synaptic signaling | 2.59E-05 | 264 | 15 |
| GO:0099537 | trans-synaptic signaling | 3.01E-05 | 267 | 15 |
| GO:0099536 | synaptic signaling | 5.42E-05 | 279 | 15 |
| GO:0050804 | modulation of chemical synaptic transmission | 0.0008816 | 175 | 11 |
| GO:0099177 | regulation of trans-synaptic signaling | 0.0009332 | 176 | 11 |
| GO:0051049 | regulation of transport | 0.0033332 | 726 | 21 |
| GO:0006810 | transport | 0.0076065 | 1879 | 36 |
| GO:1901021 | positive regulation of calcium ion transmembrane transporter activity | 0.0122566 | 15 | 4 |
| GO:0098662 | inorganic cation transmembrane transport | 0.016144 | 236 | 11 |
| GO:0051234 | establishment of localization | 0.0194168 | 1955 | 36 |
| GO:0065008 | regulation of biological quality | 0.0302319 | 1741 | 33 |
| GO:0042391 | regulation of membrane potential | 0.0355326 | 168 | 9 |
| GO:0032879 | regulation of localization | 0.0395402 | 1210 | 26 |
| GO:0098660 | inorganic ion transmembrane transport | 0.044306 | 263 | 11 |

### Transcriptional upregulation of immune response-related genes occurs after deafening

The GSEA results definitively show that a large fraction of the genes being upregulated after deafening are related to an inflammatory/immune response. Moreover, when using the top 500 most significant genes that had increased expression after deafening as input to gProfiler, most of the top enriched terms were immune response-related (Figure 1C, Table 4). These included categories such as “immune system process”, “positive regulation of immune system process”, “leukocyte activation”, and several others. We first attempted to distinguish innate or adaptive elements of the immune response by generating heatmaps of genes under the GO categories GO:0045087 – innate immune response (Figure 5A, 554 genes) or GO:0002250 – adaptive immune response (Figure 5B, 344 genes). As indicated by the heatmaps, there were several genes in both categories with increased expression after deafening, indicating that both innate and adaptive immune responses are evident after deafening. However, considering only these numbers is insufficient to determine the relative contribution of the innate or adaptive response, as there are many genes (145 of the genes in this dataset) that are present in both GO categories. We therefore focused on expression of more specific gene sets, including chemokines and chemokine receptors (Figure 6A), interleukin genes (Figure 6B) antigen presentation (Figure 6C), and complement genes (Figure 6D). As shown in Figure 6A, while many chemokines were expressed at very low levels or did not change in expression after deafening, there a few that markedly increased in expression. These included Ccl2, Cxcl10, Cxcl11, and Cxcl14, all of which were significantly differentially expressed. Ccl2, Cxcl10, and Cxcl11 had log_2_FCs>1 in deafened compared to hearing at P60 and/or P32. As these chemokines are known to recruit and activate macrophages and other immune cells including T cells (Carr et al., 1994; Cole et al., 1998; Dufour et al., 2002), it is possible that these are the major chemokines responsible for recruiting immune cells to the spiral ganglion after deafening. Ccl2, Cxcl10, and Cxcl11 are IFN-γ-inducible genes, as are many genes upregulated postdeafening. While this implies elevated IFN-γ in the cochlea, we unexpectedly did not detect significant IFN-γ expression in the ganglion in our RNAseq data. Possibly, the IFN-γ is being produced in adjacent tissues outside of the spiral ganglion.

**Figure 5.**
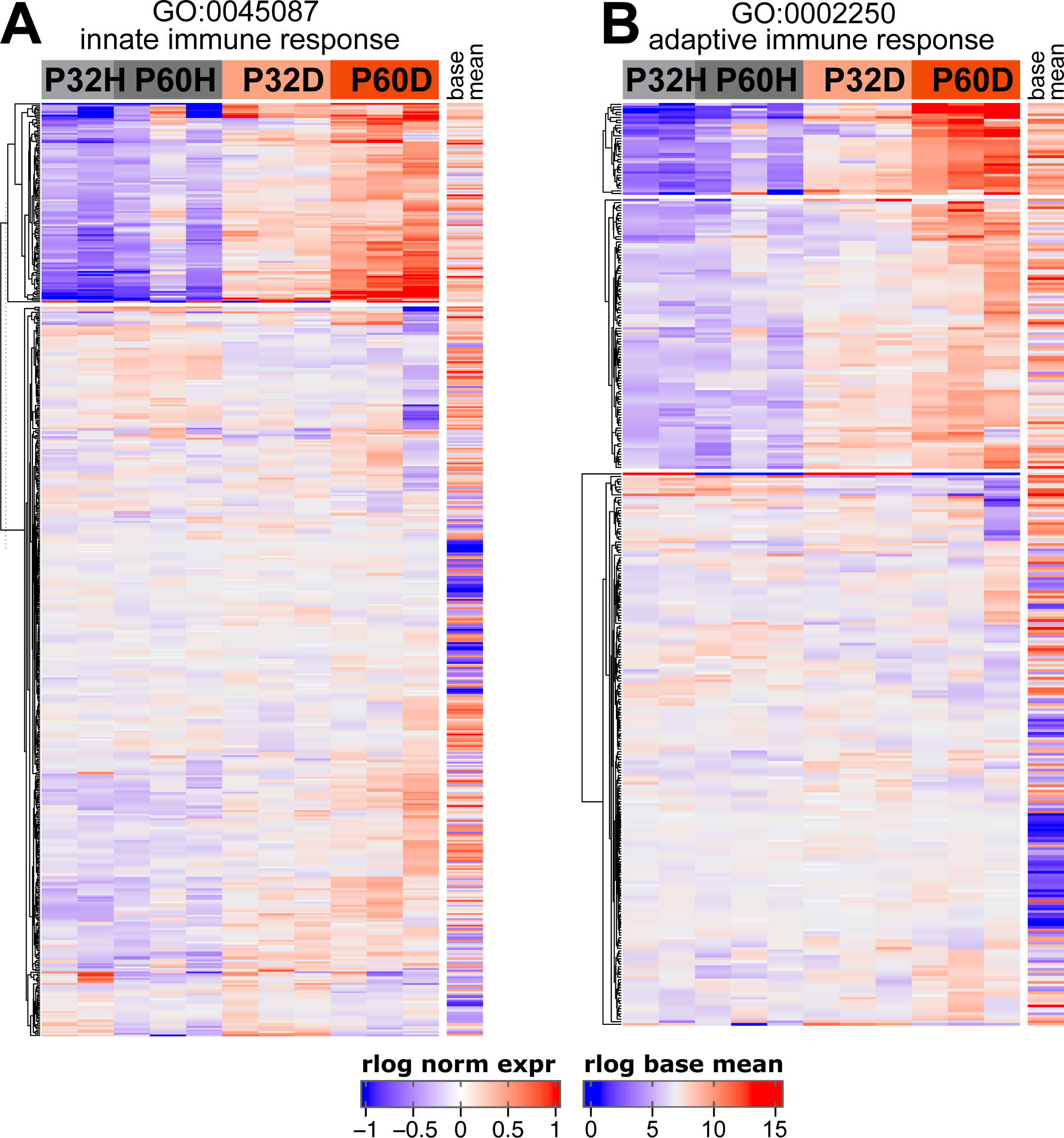
Both innate and adaptive immune response-related genes are upregulated after deafening. A) Heatmap showing regularized logarithm (rlog) normalized expression of genes under the GO term GO:0045087 – innate immune response. B) Heatmap of genes under the GO term GO:0002250 – adaptive immune response. Base mean = averaged expression over all conditions/replicates.

**Figure 6.**
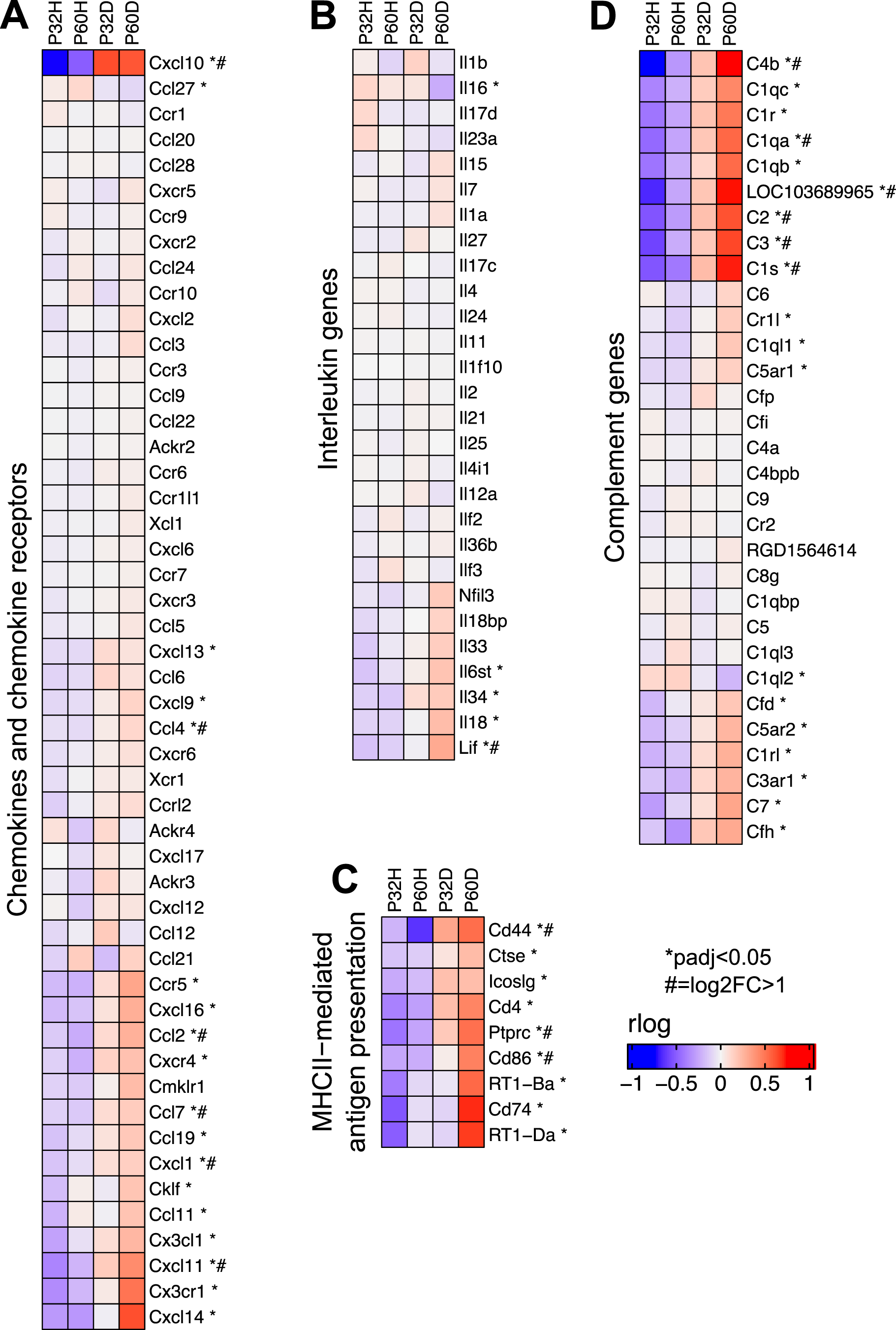
Specific groups of immune response genes upregulated after deafening. A) Heatmap showing expression of chemokine and chemokine receptor genes. B) Heatmap showing expression of interleukin genes. C) Heatmap showing expression of genes involved in MHCII-mediated antigen presentation. D) Heatmap showing expression of complement and complement receptor genes. All heatmaps show regularized logarithm (rlog) normalized expression. The expression shown for each condition is the averaged normalized expression of the biological replicates for that condition.

**Table 4.** Genes with higher expression at P60 in deafened relative to hearing (P60D/P60H). GO terms associated with inflammatory/immune responses are highlighted.

| GO term ID | GO term name | p value | term size | intersection |
| --- | --- | --- | --- | --- |
| GO:0002376 | immune system process | 3.14E-13 | 1513 | 56 |
| GO:0002684 | positive regulation of immune system process | 4.88E-13 | 533 | 34 |
| GO:0051239 | regulation of multicellular organismal process | 1.79E-12 | 1305 | 51 |
| GO:0001775 | cell activation | 2.81E-12 | 602 | 35 |
| GO:0051240 | positive regulation of multicellular organismal process | 5.69E-12 | 735 | 38 |
| GO:0002682 | regulation of immune system process | 1.31E-11 | 754 | 38 |
| GO:0045321 | leukocyte activation | 6.44E-11 | 551 | 32 |
| GO:0002252 | immune effector process | 4.64E-10 | 376 | 26 |
| GO:0019882 | antigen processing and presentation | 4.77E-10 | 77 | 14 |
| GO:0050776 | regulation of immune response | 8.20E-10 | 454 | 28 |
| GO:0048002 | antigen processing and presentation of peptide antigen | 1.58E-09 | 53 | 12 |
| GO:0070887 | cellular response to chemical stimulus | 3.12E-09 | 1400 | 48 |
| GO:0048584 | positive regulation of response to stimulus | 3.67E-09 | 1093 | 42 |
| GO:0001817 | regulation of cytokine production | 4.96E-09 | 349 | 24 |
| GO:0006955 | immune response | 5.05E-09 | 1005 | 40 |
| GO:0002274 | myeloid leukocyte activation | 5.98E-09 | 111 | 15 |
| GO:0001816 | cytokine production | 6.73E-09 | 354 | 24 |
| GO:0051094 | positive regulation of developmental process | 9.37E-09 | 661 | 32 |
| GO:0002697 | regulation of immune effector process | 4.60E-08 | 175 | 17 |
| GO:2000026 | regulation of multicellular organismal development | 4.97E-08 | 661 | 31 |
| GO:0050778 | positive regulation of immune response | 6.02E-08 | 358 | 23 |
| GO:0042221 | response to chemical | 8.61E-08 | 1950 | 55 |
| GO:0002703 | regulation of leukocyte mediated immunity | 9.57E-08 | 112 | 14 |
| GO:0007166 | cell surface receptor signaling pathway | 2.15E-07 | 1291 | 43 |
| GO:0002699 | positive regulation of immune effector process | 2.19E-07 | 119 | 14 |
| GO:0032635 | interleukin-6 production | 2.71E-07 | 80 | 12 |
| GO:0032675 | regulation of interleukin-6 production | 2.71E-07 | 80 | 12 |
| GO:0050896 | response to stimulus | 4.41E-07 | 4192 | 84 |
| GO:0010033 | response to organic substance | 7.00E-07 | 1508 | 46 |
| GO:0008283 | cell population proliferation | 1.25E-06 | 992 | 36 |
| GO:0002694 | regulation of leukocyte activation | 1.38E-06 | 382 | 22 |
| GO:0022409 | positive regulation of cell-cell adhesion | 1.77E-06 | 139 | 14 |
| GO:0019883 | antigen processing and presentation of endogenous antigen | 1.94E-06 | 29 | 8 |
| GO:0002483 | antigen processing and presentation of endogenous peptide antigen | 1.94E-06 | 29 | 8 |
| GO:0042127 | regulation of cell population proliferation | 2.49E-06 | 820 | 32 |
| GO:0002443 | leukocyte mediated immunity | 2.58E-06 | 257 | 18 |
| GO:0050865 | regulation of cell activation | 3.27E-06 | 400 | 22 |
| GO:0008284 | positive regulation of cell population proliferation | 3.38E-06 | 478 | 24 |
| GO:0050793 | regulation of developmental process | 4.01E-06 | 1193 | 39 |
| GO:0001819 | positive regulation of cytokine production | 4.09E-06 | 233 | 17 |
| GO:0002822 | regulation of adaptive immune response based on somatic recombination of immune receptors built from immunoglobulin superfamily domains | 4.81E-06 | 81 | 11 |
| GO:0042098 | T cell proliferation | 4.85E-06 | 102 | 12 |
| GO:0007159 | leukocyte cell-cell adhesion | 6.69E-06 | 181 | 15 |
| GO:0051716 | cellular response to stimulus | 7.74E-06 | 3326 | 71 |
| GO:0002819 | regulation of adaptive immune response | 1.18E-05 | 88 | 11 |
| GO:0002696 | positive regulation of leukocyte activation | 1.19E-05 | 283 | 18 |
| GO:0048856 | anatomical structure development | 1.46E-05 | 2846 | 64 |
| GO:0050867 | positive regulation of cell activation | 1.49E-05 | 287 | 18 |
| GO:0098609 | cell-cell adhesion | 1.49E-05 | 358 | 20 |
| GO:0002366 | leukocyte activation involved in immune response | 1.50E-05 | 137 | 13 |
| GO:0032502 | developmental process | 1.57E-05 | 3071 | 67 |
| GO:0042110 | T cell activation | 1.70E-05 | 256 | 17 |
| GO:0070661 | leukocyte proliferation | 1.81E-05 | 166 | 14 |
| GO:0048583 | regulation of response to stimulus | 2.07E-05 | 1855 | 49 |
| GO:1903039 | positive regulation of leukocyte cell-cell adhesion | 2.14E-05 | 116 | 12 |
| GO:0002263 | cell activation involved in immune response | 2.31E-05 | 142 | 13 |
| GO:0007155 | cell adhesion | 3.14E-05 | 623 | 26 |
| GO:0048731 | system development | 3.35E-05 | 2407 | 57 |
| GO:0022610 | biological adhesion | 3.58E-05 | 627 | 26 |
| GO:0022407 | regulation of cell-cell adhesion | 3.89E-05 | 206 | 15 |
| GO:0071310 | cellular response to organic substance | 4.50E-05 | 1133 | 36 |
| GO:0002428 | antigen processing and presentation of peptide antigen via MHC class Ib | 4.65E-05 | 28 | 7 |
| GO:0009605 | response to external stimulus | 4.68E-05 | 1415 | 41 |
| GO:0002700 | regulation of production of molecular mediator of immune response | 5.15E-05 | 79 | 10 |
| GO:0050900 | leukocyte migration | 5.48E-05 | 181 | 14 |
| GO:0006954 | inflammatory response | 6.42E-05 | 280 | 17 |
| GO:0046649 | lymphocyte activation | 7.25E-05 | 474 | 22 |
| GO:0046651 | lymphocyte proliferation | 7.74E-05 | 157 | 13 |
| GO:0002475 | antigen processing and presentation via MHC class Ib | 7.84E-05 | 30 | 7 |
| GO:1903037 | regulation of leukocyte cell-cell adhesion | 8.35E-05 | 158 | 13 |
| GO:0032943 | mononuclear cell proliferation | 9.00E-05 | 159 | 13 |
| GO:0016477 | cell migration | 9.59E-05 | 705 | 27 |
| GO:0032680 | regulation of tumor necrosis factor production | 0.000105 | 85 | 10 |
| GO:0032640 | tumor necrosis factor production | 0.000105 | 85 | 10 |
| GO:0071706 | tumor necrosis factor superfamily cytokine production | 0.000118 | 86 | 10 |
| GO:1903555 | regulation of tumor necrosis factor superfamily cytokine production | 0.000118 | 86 | 10 |
| GO:0002718 | regulation of cytokine production involved in immune response | 0.000118 | 47 | 8 |
| GO:0002367 | cytokine production involved in immune response | 0.000118 | 47 | 8 |
| GO:0002275 | myeloid cell activation involved in immune response | 0.000118 | 47 | 8 |
| GO:0050870 | positive regulation of T cell activation | 0.000126 | 110 | 11 |
| GO:0097529 | myeloid leukocyte migration | 0.000138 | 111 | 11 |
| GO:0002705 | positive regulation of leukocyte mediated immunity | 0.000151 | 67 | 9 |
| GO:0030154 | cell differentiation | 0.000155 | 1970 | 49 |
| GO:0045785 | positive regulation of cell adhesion | 0.000169 | 230 | 15 |
| GO:0071216 | cellular response to biotic stimulus | 0.000192 | 141 | 12 |
| GO:0002478 | antigen processing and presentation of exogenous peptide antigen | 0.000195 | 21 | 6 |
| GO:0031343 | positive regulation of cell killing | 0.000199 | 34 | 7 |
| GO:1902107 | positive regulation of leukocyte differentiation | 0.000203 | 91 | 10 |
| GO:1903708 | positive regulation of hemopoiesis | 0.000203 | 91 | 10 |
| GO:0048518 | positive regulation of biological process | 0.000245 | 3116 | 65 |
| GO:0032653 | regulation of interleukin-10 production | 0.000246 | 35 | 7 |
| GO:0032613 | interleukin-10 production | 0.000246 | 35 | 7 |
| GO:0009607 | response to biotic stimulus | 0.000259 | 789 | 28 |
| GO:0048869 | cellular developmental process | 0.000288 | 2008 | 49 |
| GO:0045597 | positive regulation of cell differentiation | 0.000348 | 433 | 20 |
| GO:0019884 | antigen processing and presentation of exogenous antigen | 0.000356 | 23 | 6 |
| GO:0010628 | positive regulation of gene expression | 0.000363 | 564 | 23 |
| GO:0002709 | regulation of T cell mediated immunity | 0.000369 | 37 | 7 |
| GO:0002440 | production of molecular mediator of immune response | 0.000374 | 97 | 10 |
| GO:0098581 | detection of external biotic stimulus | 0.000423 | 13 | 5 |
| GO:0032642 | regulation of chemokine production | 0.000448 | 38 | 7 |
| GO:0007275 | multicellular organism development | 0.000461 | 2581 | 57 |
| GO:0050764 | regulation of phagocytosis | 0.000488 | 56 | 8 |
| GO:0051050 | positive regulation of transport | 0.000491 | 401 | 19 |
| GO:0002706 | regulation of lymphocyte mediated immunity | 0.000515 | 77 | 9 |
| GO:0032602 | chemokine production | 0.000541 | 39 | 7 |
| GO:0002520 | immune system development | 0.000546 | 532 | 22 |
| GO:0071345 | cellular response to cytokine stimulus | 0.00055 | 404 | 19 |
| GO:0044419 | biological process involved in interspecies interaction between organisms | 0.000561 | 819 | 28 |
| GO:0045595 | regulation of cell differentiation | 0.000593 | 771 | 27 |
| GO:0034097 | response to cytokine | 0.000601 | 448 | 20 |
| GO:0042129 | regulation of T cell proliferation | 0.000644 | 79 | 9 |
| GO:0002824 | positive regulation of adaptive immune response based on somatic recombination of immune receptors built from immunoglobulin superfamily domains | 0.000645 | 58 | 8 |
| GO:0042554 | superoxide anion generation | 0.000652 | 14 | 5 |
| GO:0002521 | leukocyte differentiation | 0.000673 | 292 | 16 |
| GO:0002821 | positive regulation of adaptive immune response | 0.000739 | 59 | 8 |
| GO:0051249 | regulation of lymphocyte activation | 0.000844 | 335 | 17 |
| GO:0042116 | macrophage activation | 0.00092 | 42 | 7 |
| GO:0009595 | detection of biotic stimulus | 0.000968 | 15 | 5 |
| GO:0002476 | antigen processing and presentation of endogenous peptide antigen via MHC class Ib | 0.001002 | 27 | 6 |
| GO:0051674 | localization of cell | 0.001089 | 795 | 27 |
| GO:0048870 | cell motility | 0.001089 | 795 | 27 |
| GO:0050863 | regulation of T cell activation | 0.001134 | 166 | 12 |
| GO:0032501 | multicellular organismal process | 0.001151 | 3622 | 70 |
| GO:0002724 | regulation of T cell cytokine production | 0.001395 | 16 | 5 |
| GO:0071276 | cellular response to cadmium ion | 0.001395 | 16 | 5 |
| GO:0002369 | T cell cytokine production | 0.001395 | 16 | 5 |
| GO:0050670 | regulation of lymphocyte proliferation | 0.001442 | 112 | 10 |
| GO:0006950 | response to stress | 0.001694 | 1797 | 44 |
| GO:0032944 | regulation of mononuclear cell proliferation | 0.001699 | 114 | 10 |
| GO:0030155 | regulation of cell adhesion | 0.001818 | 354 | 17 |
| GO:0002237 | response to molecule of bacterial origin | 0.001916 | 207 | 13 |
| GO:1903706 | regulation of hemopoiesis | 0.002022 | 208 | 13 |
| GO:0006952 | defense response | 0.002153 | 772 | 26 |
| GO:0070663 | regulation of leukocyte proliferation | 0.002159 | 117 | 10 |
| GO:0051707 | response to other organism | 0.002206 | 773 | 26 |
| GO:0048522 | positive regulation of cellular process | 0.002256 | 2842 | 59 |
| GO:0060627 | regulation of vesicle-mediated transport | 0.002267 | 245 | 14 |
| GO:0043207 | response to external biotic stimulus | 0.002316 | 775 | 26 |
| GO:0032755 | positive regulation of interleukin-6 production | 0.002364 | 48 | 7 |
| GO:0150062 | complement-mediated synapse pruning | 0.002509 | 3 | 3 |
| GO:0002495 | antigen processing and presentation of peptide antigen via MHC class II | 0.002683 | 18 | 5 |
| GO:0065009 | regulation of molecular function | 0.002711 | 1160 | 33 |
| GO:0032760 | positive regulation of tumor necrosis factor production | 0.00273 | 49 | 7 |
| GO:1903557 | positive regulation of tumor necrosis factor superfamily cytokine production | 0.00273 | 49 | 7 |
| GO:0030099 | myeloid cell differentiation | 0.002783 | 214 | 13 |
| GO:0032649 | regulation of interferon-gamma production | 0.002813 | 70 | 8 |
| GO:0032609 | interferon-gamma production | 0.002813 | 70 | 8 |
| GO:0002720 | positive regulation of cytokine production involved in immune response | 0.002919 | 32 | 6 |
| GO:0006909 | phagocytosis | 0.003088 | 216 | 13 |
| GO:1990266 | neutrophil migration | 0.003138 | 71 | 8 |
| GO:0002456 | T cell mediated immunity | 0.003143 | 50 | 7 |
| GO:0031341 | regulation of cell killing | 0.003143 | 50 | 7 |
| GO:0001818 | negative regulation of cytokine production | 0.003169 | 122 | 10 |
| GO:0002460 | adaptive immune response based on somatic recombination of immune receptors built from immunoglobulin superfamily domains | 0.003422 | 218 | 13 |
| GO:0042119 | neutrophil activation | 0.003605 | 19 | 5 |
| GO:0032930 | positive regulation of superoxide anion generation | 0.003606 | 9 | 4 |
| GO:0044093 | positive regulation of molecular function | 0.003757 | 694 | 24 |
| GO:0048534 | hematopoietic or lymphoid organ development | 0.003797 | 504 | 20 |
| GO:0023052 | signaling | 0.003799 | 2738 | 57 |
| GO:0030335 | positive regulation of cell migration | 0.00381 | 256 | 14 |
| GO:0002444 | myeloid leukocyte mediated immunity | 0.004125 | 52 | 7 |
| GO:0046686 | response to cadmium ion | 0.004246 | 34 | 6 |
| GO:0071219 | cellular response to molecule of bacterial origin | 0.004252 | 126 | 10 |
| GO:1902105 | regulation of leukocyte differentiation | 0.004274 | 156 | 11 |
| GO:0001774 | microglial cell activation | 0.00476 | 20 | 5 |
| GO:0002504 | antigen processing and presentation of peptide or polysaccharide antigen via MHC class II | 0.00476 | 20 | 5 |
| GO:0036230 | granulocyte activation | 0.00476 | 20 | 5 |
| GO:0030097 | hemopoiesis | 0.005172 | 469 | 19 |
| GO:2000147 | positive regulation of cell motility | 0.005462 | 264 | 14 |
| GO:0009617 | response to bacterium | 0.005678 | 472 | 19 |
| GO:0002708 | positive regulation of lymphocyte mediated immunity | 0.00607 | 55 | 7 |
| GO:0040011 | locomotion | 0.006075 | 869 | 27 |
| GO:0007154 | cell communication | 0.006271 | 2778 | 57 |
| GO:0007165 | signal transduction | 0.006427 | 2565 | 54 |
| GO:0033993 | response to lipid | 0.006667 | 523 | 20 |
| GO:0051272 | positive regulation of cellular component movement | 0.006794 | 269 | 14 |
| GO:0002702 | positive regulation of production of molecular mediator of immune response | 0.006867 | 56 | 7 |
| GO:0040017 | positive regulation of locomotion | 0.007403 | 271 | 14 |
| GO:0002250 | adaptive immune response | 0.008601 | 237 | 13 |
| GO:0030593 | neutrophil chemotaxis | 0.008725 | 58 | 7 |
| GO:0002474 | antigen processing and presentation of peptide antigen via MHC class I | 0.009265 | 11 | 4 |
| GO:2001185 | regulation of CD8-positive, alpha-beta T cell activation | 0.009265 | 11 | 4 |
| GO:0032928 | regulation of superoxide anion generation | 0.009265 | 11 | 4 |
| GO:2000427 | positive regulation of apoptotic cell clearance | 0.009945 | 4 | 3 |
| GO:0098883 | synapse pruning | 0.009945 | 4 | 3 |
| GO:0030595 | leukocyte chemotaxis | 0.011397 | 111 | 9 |
| GO:0048513 | animal organ development | 0.011509 | 1860 | 43 |
| GO:0097530 | granulocyte migration | 0.01231 | 85 | 8 |
| GO:0018212 | peptidyl-tyrosine modification | 0.012444 | 142 | 10 |
| GO:0018108 | peptidyl-tyrosine phosphorylation | 0.012444 | 142 | 10 |
| GO:0032735 | positive regulation of interleukin-12 production | 0.012547 | 24 | 5 |
| GO:0051241 | negative regulation of multicellular organismal process | 0.012691 | 499 | 19 |
| GO:0048143 | astrocyte activation | 0.013766 | 12 | 4 |
| GO:0002573 | myeloid leukocyte differentiation | 0.014179 | 114 | 9 |
| GO:0002698 | negative regulation of immune effector process | 0.015265 | 63 | 7 |
| GO:0071887 | leukocyte apoptotic process | 0.015265 | 63 | 7 |
| GO:0002253 | activation of immune response | 0.015342 | 250 | 13 |
| GO:0006801 | superoxide metabolic process | 0.015531 | 25 | 5 |
| GO:0002711 | positive regulation of T cell mediated immunity | 0.015531 | 25 | 5 |
| GO:0071248 | cellular response to metal ion | 0.015948 | 88 | 8 |
| GO:0002683 | negative regulation of immune system process | 0.016815 | 215 | 12 |
| GO:0043299 | leukocyte degranulation | 0.017594 | 43 | 6 |
| GO:0050766 | positive regulation of phagocytosis | 0.017594 | 43 | 6 |
| GO:0051251 | positive regulation of lymphocyte activation | 0.018193 | 254 | 13 |
| GO:0045637 | regulation of myeloid cell differentiation | 0.018775 | 118 | 9 |
| GO:0002283 | neutrophil activation involved in immune response | 0.019697 | 13 | 4 |
| GO:0055094 | response to lipoprotein particle | 0.019697 | 13 | 4 |
| GO:0019886 | antigen processing and presentation of exogenous peptide antigen via MHC class II | 0.019697 | 13 | 4 |
| GO:0080164 | regulation of nitric oxide metabolic process | 0.023142 | 27 | 5 |
| GO:0045428 | regulation of nitric oxide biosynthetic process | 0.023142 | 27 | 5 |
| GO:0010038 | response to metal ion | 0.024277 | 187 | 11 |
| GO:2000425 | regulation of apoptotic cell clearance | 0.02464 | 5 | 3 |
| GO:0090322 | regulation of superoxide metabolic process | 0.027315 | 14 | 4 |
| GO:0001916 | positive regulation of T cell mediated cytotoxicity | 0.027315 | 14 | 4 |
| GO:0016192 | vesicle-mediated transport | 0.02987 | 731 | 23 |
| GO:1903131 | mononuclear cell differentiation | 0.030312 | 228 | 12 |
| GO:0019221 | cytokine-mediated signaling pathway | 0.03262 | 193 | 11 |
| GO:0001912 | positive regulation of leukocyte mediated cytotoxicity | 0.033381 | 29 | 5 |
| GO:0050798 | activated T cell proliferation | 0.033381 | 29 | 5 |
| GO:0002704 | negative regulation of leukocyte mediated immunity | 0.033381 | 29 | 5 |
| GO:0071621 | granulocyte chemotaxis | 0.033846 | 71 | 7 |
| GO:0071402 | cellular response to lipoprotein particle stimulus | 0.036895 | 15 | 4 |
| GO:0021782 | glial cell development | 0.038011 | 49 | 6 |
| GO:0032496 | response to lipopolysaccharide | 0.03947 | 197 | 11 |
| GO:0006809 | nitric oxide biosynthetic process | 0.039667 | 30 | 5 |
| GO:2001057 | reactive nitrogen species metabolic process | 0.046835 | 31 | 5 |
| GO:0046209 | nitric oxide metabolic process | 0.046835 | 31 | 5 |
| GO:0042102 | positive regulation of T cell proliferation | 0.047987 | 51 | 6 |
| GO:0032620 | interleukin-17 production | 0.048729 | 16 | 4 |
| GO:0001914 | regulation of T cell mediated cytotoxicity | 0.048729 | 16 | 4 |
| GO:0032660 | regulation of interleukin-17 production | 0.048729 | 16 | 4 |
| GO:0032490 | detection of molecule of bacterial origin | 0.048839 | 6 | 3 |
| GO:0006928 | movement of cell or subcellular component | 0.049164 | 974 | 27 |

Interleukin molecules, such as IL-1β, IL-6, and IL-18 are potent inflammatory mediators that are often involved in neuroinflammatory and neurodegenerative processes (Felderhoff-Mueser et al., 2005; Smith et al., 2012; Kummer et al., 2021). As shown in Figure 6B, IL-18 is expressed in the spiral ganglion although its expression is not significantly elevated by deafening. In contrast, IL-6 was not detected in our dataset and most interleukins, including IL-1β, were expressed at low levels (base mean < 100); few exhibited changes in expression after deafening (Figure 6C). These data imply that interleukin signaling plays only a small role in induction of the post-deafening immune response and neurodegeneration, although pro-inflammatory interleukins that are expressed, e.g. IL-18, may contribute to the response in a permissive manner.

Key genes for local induction of the adaptive immune response are those involved in antigen processing and presentation. As shown in Figure 6C several of these genes have significantly increased expression after deafening at P60 and/or P32. These included MHCII components RT1-Da and RT1-Ba, Cd74 (MHCII invariant chain), Ctse (cathepsin e), Cd4, Icoslg and Cd86 (MHCII co-stimulatory signaling molecules), Cd44 and Cd69 (T cell activation markers). These genes are all required for antigen presentation, typically from macrophages to T cells. Their increased expression suggests that this pathway is activated in the spiral ganglion after deafening and raises the possibility that antigen presentation and the adaptive immune response could be contributing to SGN death.

### Upregulation of complement cascade genes occurs after deafening

Among immune response-related genes significantly upregulated after deafening were genes encoding early complement pathway components C1-C4 (Figure 6D). These were elevated post-deafening at P32 and at P60, indicating that complement activation is initiated early in the response to deafening and is persistent. Genes for the late complement components (C5-C9), which contribute to membrane attack complex formation, are mainly synthesized in the liver (Lubbers et al., 2017) so, other than a modest elevation in C7, did not change in expression after deafening. Finally, several of the complement receptors, including C3ar1, C5ar1/2, Cd11b, and Cd18, were upregulated after deafening, indicating that cells capable of responding to complement are present and are either increasing in number or increasing responsiveness after deafening. These results imply that complement activation could be contributing to the deafening-related immune response and subsequent SGN death. Because C3 is involved in all complement activation pathways and is the main mediator in complement activation (Noris and Remuzzi, 2013), we chose to assess SGN survival and immune response activation after deafening in C3 knockout rats.

### C3 knockout does not affect normal hearing or aminoglycoside-induced hair cell loss

Because complement, and, specifically, C3, is known to be involved in synapse pruning in and neural circuit development in the CNS (Schafer et al., 2012) and retina (Anderson et al., 2019), we assessed hearing function in the C3KO rats using ABR measurements. As described in Materials and Methods, normal hearing WT, heterozygous, and C3KO rats underwent ABR testing at P32 to measure ABR thresholds, wave I amplitude, and wave I latency at 4, 8, and 16 kHz. Representative ABR waveforms from individual mice in response to 8 kHz suprathreshold stimuli at are shown in Supplemental Figure 4A. ABR thresholds were similar across genotype at all frequencies measured (Supplemental Figure 4B). Similarly, neither wave I amplitudes (Supplemental Figure 4D) nor latencies (Supplemental Figure 4D) differed significantly among genotypes at all frequencies measured.

We also considered the possibility that knockout of C3 might ameliorate kanamycin-induced hair cell loss. To confirm that hair cell loss did occur in kanamycin-treated C3KO rats, we viewed hair cells in cochlear wholemount preparations from hearing and deaf rats of each genotype, wild-type (C3^+/+^), heterozygous (C3^+/–^), C3 knockout (C3^−/–^).The cochleae were labeled with combined anti-myosin VIIa and anti-myosin VI to view hair cells and anti-NF200 to view neurons. Both inner and outer hair cells appeared normal in C3^−/–^ and C3^+/–^ rats compared to WT rats with no apparent missing hair cells (Supplemental Figure 5A-C). Hair cell loss was complete in kanamycin-injected rats of all three genotypes (Supplemental Figure 5D-F). Together, these results indicate that neither normal hearing nor kanamycin-induced hair cell loss are affected by loss of C3.

### C3 knockout does not affect SGN survival post-deafening

Upregulation of early complement component (C1-4) genes was observed in the ganglion of deafened rats at both P32 and P60, suggesting that complement activation is triggered after hair cell loss. Based on these data and previous studies implicating complement, specifically C3, in other neuropathies (Royer et al., 2019), we hypothesized that complement activation was, at least in part, triggering the deafening-related immune response and/or directly contributing to SGN death after hair cell loss. To test this hypothesis, we compared SGN survival in deafened WT, C3 heterozygous, and C3KO rats to age-and genotype-matched littermate controls.

Representative images of cross sections of the ganglion showing SGNs are shown in Figure 7A-C. Hearing WT animals had an average of 2,192 SGNs/mm^2^ across all cochlear locations. The density was slightly lower in the base (Figure 7A1) than in the apex (Figure 7A4). Hearing heterozygous and hearing C3KO rats had slightly but not significantly higher overall average SGN densities (images not shown). As with WT animals, both heterozygous and C3KO animals had lower SGN densities in the base than the apex. SGN loss was observed in deafened animals of genotypes and across cochlear location. The decrease in SGN density was statistically significant (p<0.0001, 2-way ANOVA). In the base of deafened animals, SGN density was 67% of hearing controls in WT rats (Figure 7B1) and 74% of controls in both heterozygous and C3KO rats (Figure 7C1). The difference in the base was only statistically significant for WT rats. SGN loss in the middle turns was greater than that in the base, with SGN density being 61–65% of normal in the first middle turn and 53–58% of normal in the second middle turn. The degree of SGN loss was not significantly different among the genotypes but was significantly different in deafened vs. hearing animals in all genotypes (p<0.001 in all cases, 2-way ANOVA with Tukey’s multiple comparisons test, Figure 7D). SGN loss in deafened vs. hearing animals was also significant in the apex (Figure 7A-C4, D) and occurred to similar degrees in all genotypes. SGN densities were 77%, 65%, and 70% of normal in deafened WT, heterozygous, and C3KO rats, respectively. These results indicate that complement activation, C3 in particular, is not involved in SGN degeneration post-deafening.

**Figure 7.**
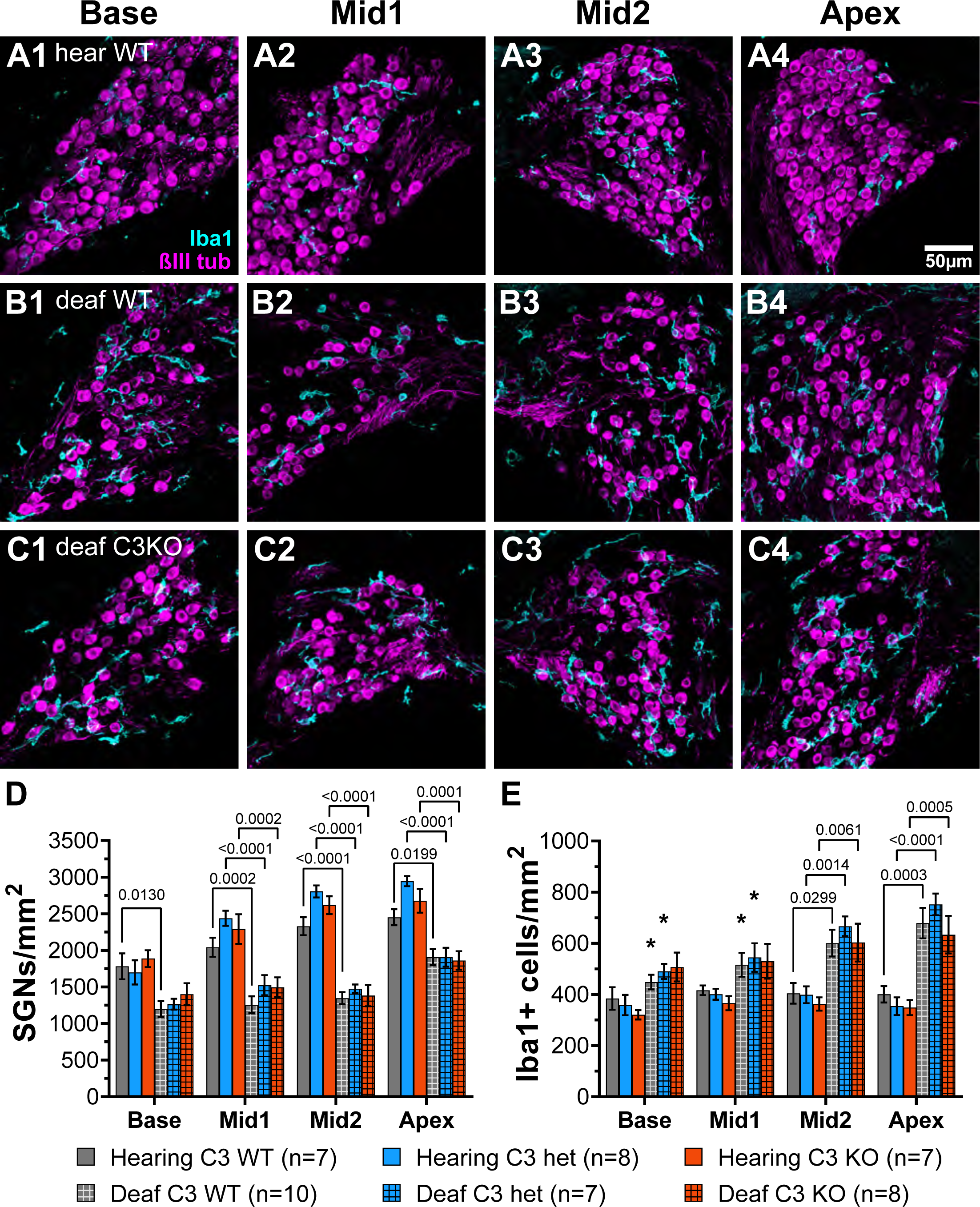
Complement is not required for SGN death or macrophage infiltration after deafening. A-C) Representative images of spiral ganglion cross-sections showing SGNs (magenta) and macrophages (cyan) in hearing WT (A1-4), deaf WT (B1-4), and deaf C3KO (C1-4) rats. Different cochlear locations are shown across the columns: base (1), middle 1 (2), middle 2 (3), and apex (4). D) Graph showing quantification of SGN density (per mm^2^) in Rosenthal’s canal. E) Quantification of macrophage density (Iba1+ cells/mm^2^) in Rosenthal’s canal at each cochlear location. The p-values were calculated using a 2-way ANOVA with Tukey’s multiple comparisons. The stars in E indicate significant differences between the indicated turns and the apex within that genotype. WT: wildtype; het: heterozygous; C3KO: C3 knockout; n = number of animals.

### C3 knockout does not affect macrophage infiltration or activation post-deafening

Though the data on SGN survival in C3KO rats suggest that C3 is not required for an immune-related contribution to neuronal death post-deafening, the possibly that C3 plays a role in recruitment and activation of macrophages in the ganglion remains. Cochlear sections used for counting neurons were also labeled with anti-Iba1 and anti-CD68 to detect, respectively, macrophages and activated macrophages (Figure 7A-C). Similar to previous observations in rats (Rahman et al., 2023) and mice (Sato et al., 2010; Kaur et al., 2015), there was a resident macrophage population present in the spiral ganglion of hearing WT rats (Figure 7A1-4). The average macrophage densities for heterozygous and C3KO rats (images not shown) were slightly lower than WT, but these differences were not statistically significant. Moreover, there were no differences in macrophage density across cochlear location in hearing rats of any of the genotypes.

There was a significant difference in macrophage density in deafened compared to hearing rats (p<0.0001, 2-way ANOVA, Figure 7E). In the base, macrophage density increased only slightly in deafened WT (Figure 7B1) compared to hearing WT (Figure 7A1) rats. Similarly, there were also only slight increases in deafened heterozygous and C3KO rats. These differences were not significant. The same was true for the basal middle segment. In the apical middle segment (Figure 7A3, B3, and C3) and apex (Figure 7A4, B4, and C4). Quantification macrophage density was significantly higher in deafened compared to hearing rats in all three genotypes (Figure 7E). We also observed differences in macrophage density in deafened animals based on cochlear location. In deafened WT animals (Figure 7B1-4), the macrophage density in the apex was significantly higher than in the base (p<0.05) and first middle turn (p<0.05). The macrophage density in the apex of deafened heterozygous animals was also significantly higher than in the base (p<0.05) and first middle turn (p<0.05). No significant differences in macrophage density were observed in deafened animals among the genotypes at any specific cochlear location.

As complement is known to play a role in macrophage activation (Dailey et al., 1998), we also quantified and compared macrophage activation among hearing and deafened animals and across genotypes by calculating the percentage of macrophages that had elevated CD68 expression (CD68+). As shown in Figure 8, the percentage of macrophages that were CD68+ increased from <10% in hearing rats to ∼50% in deafened rats. Similar increases were observed in all genotypes, and this increase in CD68+ macrophages was statistically significant for all genotypes (p<0.0001, 2-way ANOVA with Tukey’s post-hoc). In contrast with our observations that macrophage density was greater in the apex than the base, macrophage activation did not appear to vary along the base-apex axis (Figure 8B). Along with the above results on macrophage infiltration, these results indicate that complement activation is not required for macrophage infiltration and activation in the spiral ganglion of deafened rats.

**Figure 8.**
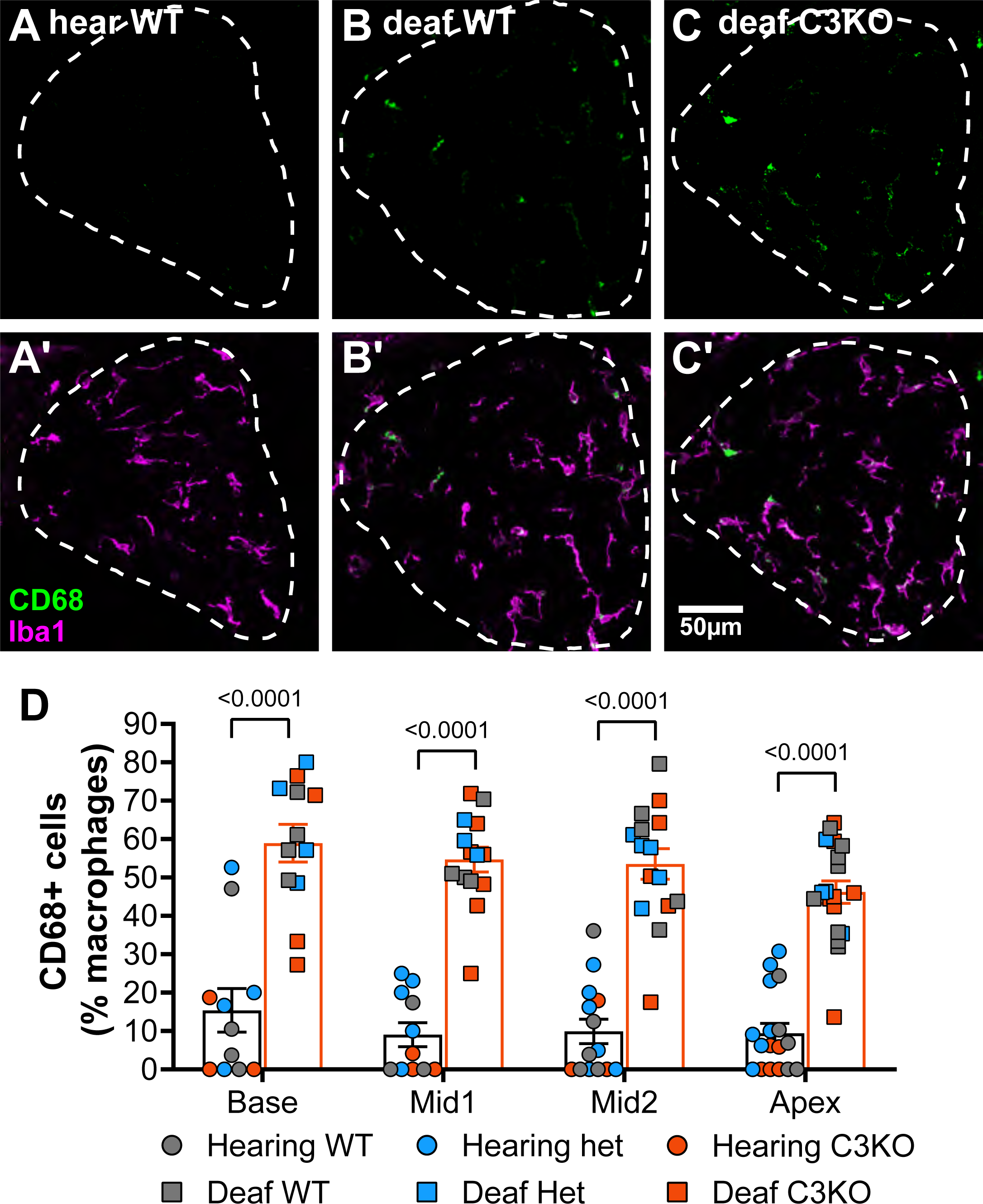
C3 is not required for macrophage activation after deafening. A-C) Representative images showing CD68 (green, A-C) alone or CD68 with Iba1 (magenta, A’-C’) in hearing WT (A-A’), deafened WT (B-B’), and deafened C3KO (C-C’) animals. D) Graph showing quantification of the percentage of macrophages that are CD68+ across cochlear location in hearing and deafened rats. Columns showing means and standard deviations are for combined all genotypes combined, with counts for different genotypes being shown as individual points. No significant differences across cochlear location or among genotypes were observed. Statistical values were determined by 2-way ANOVA with Tukey’s multiple comparisons. WT: wildtype; het: heterozygous; KO: knockout. n = number of animals.

### Gene expression is not different between male and female rats

To verify the results of the RNAseq, we selected some representative neuronal and immune system genes and quantified their expression by qPCR at P60. Although the RNAseq combined ganglia from male and female rats, for the qPCR assays male and female ganglia were separately assayed to allow detection of sex differences. For the immune-related genes (Figure 9A), we assessed expression of Cd68, C1qa, C3, Cxcl10, RT1-Da, and Cd4, as these genes covered the major groups identified in the RNAseq (activated macrophages, chemokines, complement, and MHCII-mediated antigen presentation). There were no differences in expression of immune system genes between male and female rats, whether hearing or deafened. The neuronal genes assessed were Nefh, Kcnc1, Kcnc3, Kcnd2, Scn8a, Scn4b, and Syp (Figure 9B). Similar to the immune genes, expression of these genes did not differ between males and females, except Kcnc3, which appeared to have lower expression in deafened males compared to deafened females. In agreement with the RNAseq, all immune genes assessed were upregulated in deafened compared to hearing rats, while all neuronal genes except Kcnd2 were downregulated (Figure 9C) in deafened rats. These data suggest that the post-deafening response in the spiral ganglion is similar in male and female rats.

**Figure 9.**
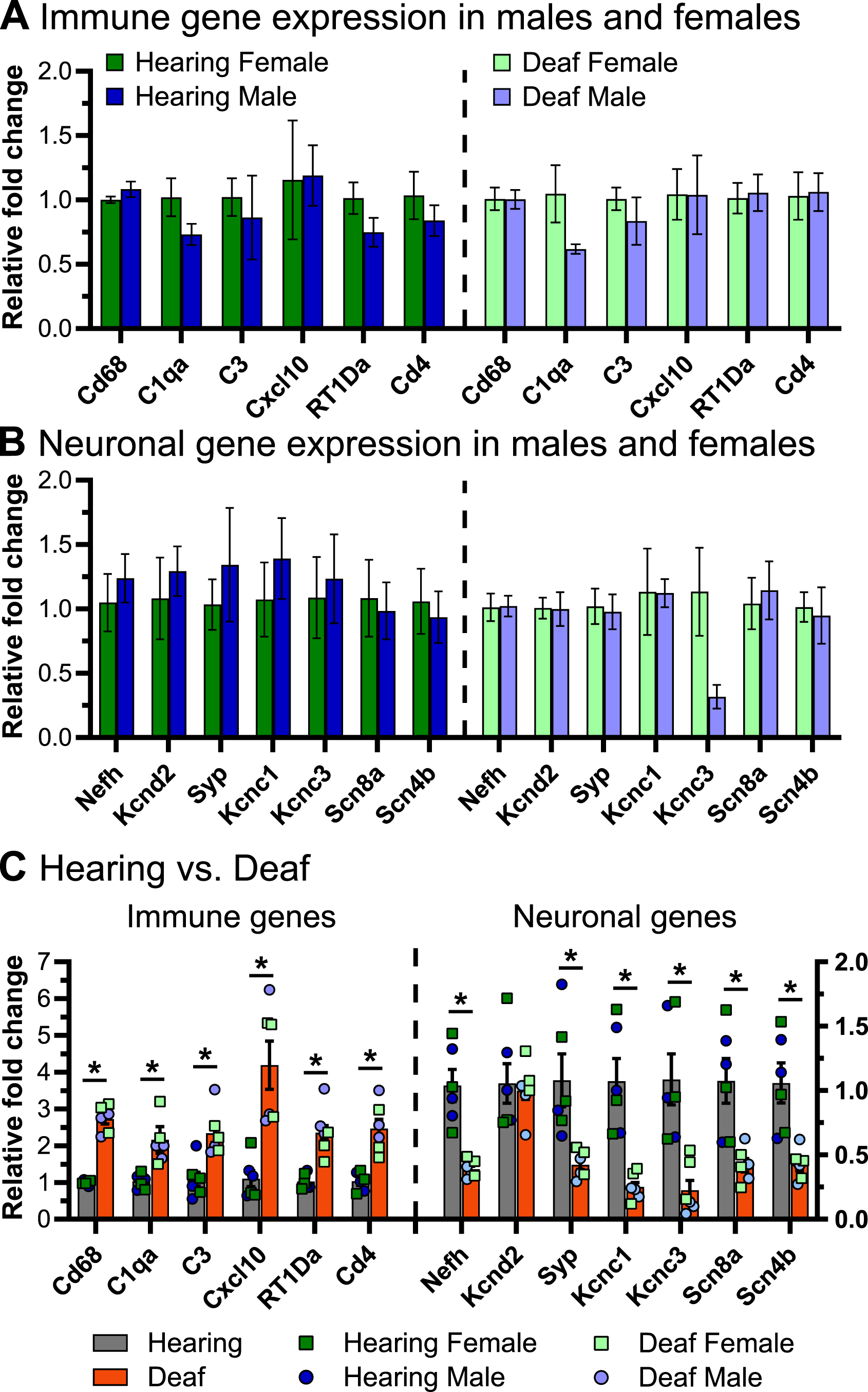
Expression of select immune-related and neuronal genes in the spiral ganglia of male and female rats. qPCR was performed as described in Materials and Methods to measure gene expression of select immune-related genes and select neuronal genes. A) Relative fold change of immune-related genes comparing males vs. females in hearing animals (left) and deafened animals (right). B) Relative fold change of neuronal genes comparing males vs. females in hearing animals (left) and deafened animals (right). Three biological replicates were used for each condition. C) Relative fold change of immune genes (left) and neuronal genes (right) comparing hearing to deafened rats. Data from male and female rats was combined, but the individual data points are shown on the graph. Immune genes are plotted on the left y-axis and neuronal genes are plotted on the right y-axis. *FDR q-value < 0.01, multiple unpaired t-tests with Benjamini, Kreiger, and Yekutieli FDR multiple comparisons. FDR: false discovery rate.

### SGN loss and increased macrophage number occur similarly in male and female rats

We further asked whether there were sex differences in numbers of SGNs and of macrophages in the spiral ganglion, comparing SGN and Iba1+ cell (macrophage) densities in both hearing and deafened rats in males and females. Because there were no observed differences among C3 genotypes, rats of all genotypes (C3 WT, heterozygous, and C3KO) were combined for this analysis. SGN and macrophage densities comparing male and female rats are shown in Figure 10. Neither SGN (Figure 10A) nor macrophage density (Figure 10B) were significantly different between male and female hearing rats at any cochlear location. Also, neither SGN (Figure 10A) nor macrophage density (Figure 10B) were significantly different between male and female deafened rats at any cochlear location. These data indicate that not only are numbers of neurons and macrophages similar in the spiral ganglia of male and female rats but also that male and female rats exhibit quantitatively similar neuronal loss and increased macrophage density after deafening.

**Figure 10.**
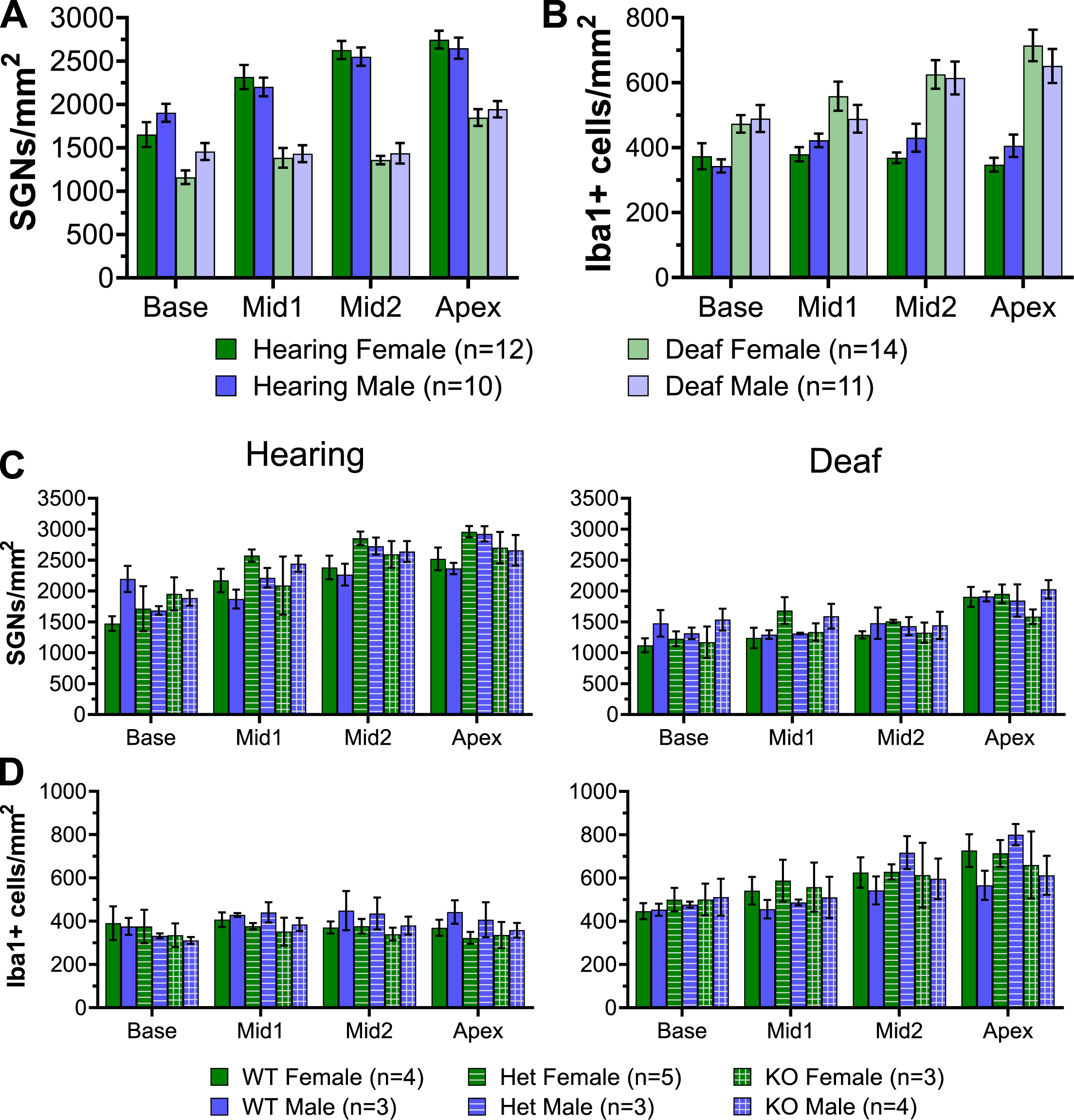
SGN and macrophage densities in the spiral ganglia of male and female rats. A) Graph showing quantification of SGN density in hearing and deafened female (green) and male (blue) rats. All genotypes were combined. B) Graph showing quantification of Iba1+ cell density in hearing and deafened female and male rats. All genotypes were combined. C) Graph showing quantification of SGN density in hearing (left) and deafened (right) female and male rats, separated by genotype. D) Graph showing quantification of Iba1+ cell density in hearing (left) and deafened (right) female and male rats, separated by genotype. No significant differences were observed between males and females for any comparison. n = number of animals.

### Lymphocytes infiltrate the spiral ganglion after deafening

The RNAseq showed that upregulation of immune-related genes after deafening is not limited to the innate immune response but also includes many genes associated with the adaptive immune response (Figure 5), which involves lymphocytes such as T cells and B cells. For example, as shown in Figure 5, genes associated with antigen processing and presentation, such as MHCII molecules (e.g., RT1-Da), lymphocyte markers (e.g., Ptprc/CD45), and T cell markers (e.g., Cd4, Cd44, and Cd86) are upregulated after deafening. To confirm involvement of cells associated with the adaptive immune system, we labeled cochlear sections with an antibody to CD45, a general immune cell marker that has been previously used for immunohistochemistry in rat (Wu et al., 1997; Metzger et al., 2002). This should label all immune cells, including macrophages and lymphocytes. As expected, macrophages in the ganglion labeled positively for CD45 (Figure 11A). Macrophages also label positively with Iba1. To distinguish CD 45+ lymphocytes from macrophages, the Iba1 signal was subtracted from the CD45 signal as described in Materials and Methods. This revealed CD45+/Iba1-cells that, significantly, exhibited a typical lymphocyte morphology – small, round, and with relatively large nuclei – very distinct from macrophages (Figure 11A,B). These lymphocytes were present in the hearing ganglion at low densities in all cochlear turns (Figure 11E). After deafening, the density of lymphocytes increased in all cochlear regions, with the increase being significantly greater (Mann-Whitney U test) in the middle region (Figure 11E).

**Figure 11.**
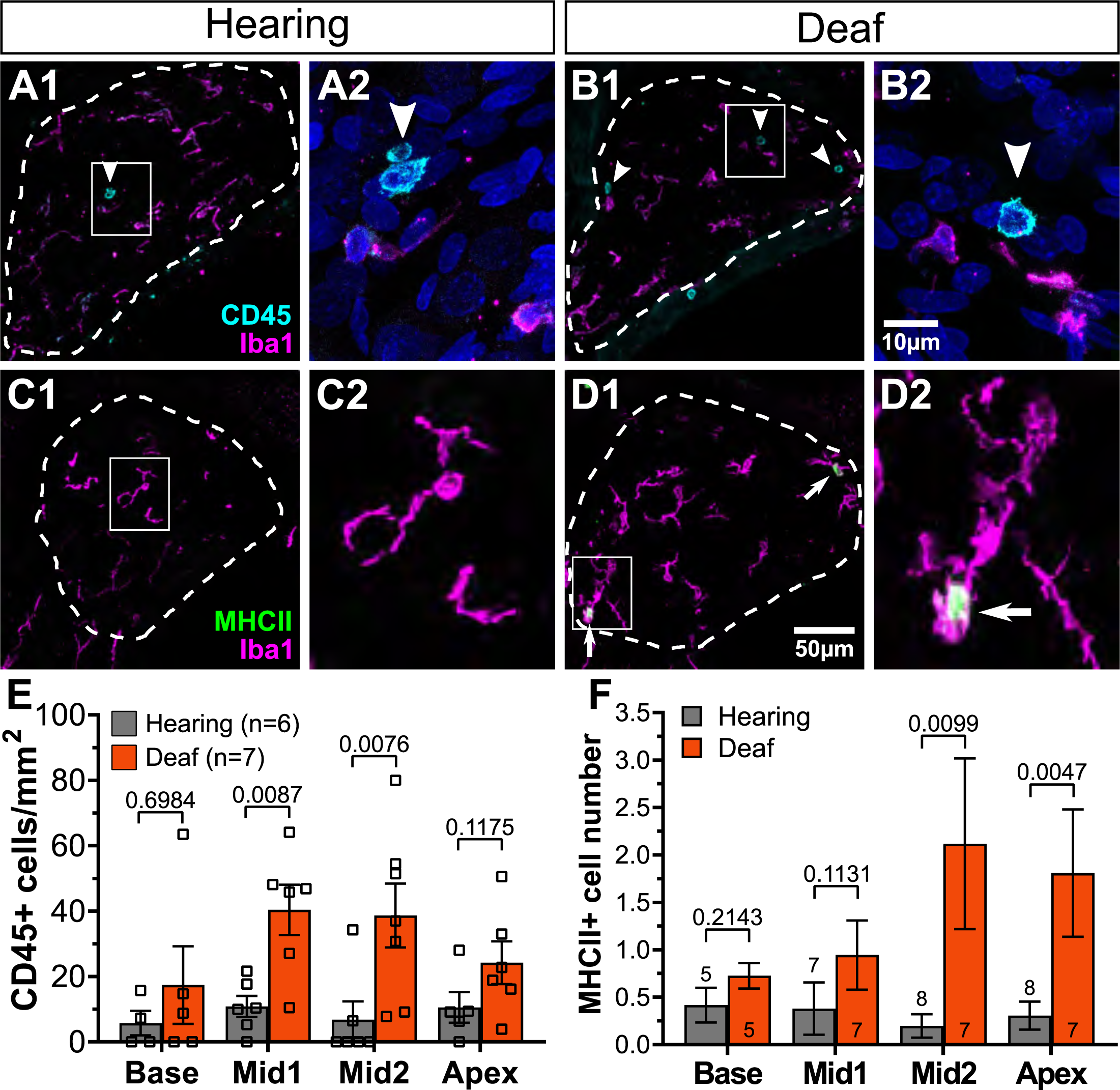
Lymphocytes and MHCII-expressing macrophages infiltrate the spiral ganglion after deafening. A-B) Representative images of CD45+ cells (cyan) and Iba1+ cells (magenta) in the hearing (A1-2) and deafened (B1-2) ganglion. The boxed areas in A1 and B1 indicated the areas of magnification shown in A2 and B2, respectively. C-D) Representative images of MHCII+ cells (green) and Iba1+ cells (magenta) in the hearing (C1-2) and deafened (D1-2) ganglion. The boxed areas in C1 and D1 indicate the areas of magnification shown in C2 and D2, respectively. E) Quantification of CD45+/Iba1-cells across cochlear location. F) Quantification of MHCII+ cells across cochlear location. n = number of animals, p-values determined Mann-Whitney U test comparing hearing and deafened at each cochlear location.

The presence of activated macrophages (Iba+/CD68+) and of lymphocytes (CD45+/Iba-) in the ganglion raises the possibility of antigen presentation by macrophages to T-lymphocytes. This is further suggested by upregulated expression of genes associated with antigen presentation (Figure 6D). We therefore asked whether MHCII-expressing cells were indeed present in the ganglion and if they were more prevalent after deafening. By combining immunolabeling with Iba1 and MHCII, we confirmed the presence of MHCII-expressing cells in the spiral ganglion and that the MHCII-expressing cells are Iba1+ macrophages (Figure 11C-D). Consistent with the post-deafening upregulation of MHCII genes observed in the RNAseq, the number of MHCII+ cells is increased after deafening (Figure 11F) with the increase significant (Mann-Whitney U test) in the apical half of the ganglia. Although the number of CD45+ lymphocytes and the number of MHCII+ cells are low relative to the number of macrophages, the increased presence of CD45+ lymphocytes and MHCII+ macrophages in the deafened ganglion implies that an adaptive immune response recruited by MHCII-dependent antigen presentation may be contributing to SGN death post-deafening.

## Discussion

Aminoglycoside-induced hair cell loss causes gradual degeneration of SGNs. This study builds on previous evidence (Hirose et al., 2005; Hirose et al., 2014; Kaur et al., 2015; Kaur et al., 2018; Rahman et al., 2023) that an immune/inflammatory response contributes to cochlear neuropathologies after cochlear trauma. We observed an increase in macrophage density and activation in the spiral ganglion post-deafening, confirming previous studies in mice (Kaur et al., 2018) and rats (Rahman et al., 2023). In addition, we observed an increase in expression of genes characteristic of inflammatory and immune responses, genes typically expressed in macrophages and lymphocytes. Increased numbers of lymphocytes in the spiral ganglion post-deafening is confirmed by direct histological observation, which also shows increased numbers of MHCII-expressing macrophages. These results, together with our previous report showing that anti-inflammatory agents reduce SGN death (Rahman et al., 2023), support hypotheses that inflammatory and immune responses occur after hair cell loss, that they contribute significantly to neurodegeneration and that both innate and adaptive immune responses are involved.

Neither normal SGN density nor SGN degeneration after deafening appeared to differ between males and females (Figure 9A,C). While this appears to contrast with a report of sex differences in responses to noise-induced damage (Rouse et al., 2020), an important difference is that we deafen rats in the second postnatal week, prior to sexual maturity, and SGN pathologies post-deafening are long-term, compared to the more acute pathologies caused by noise-induced damage.

### Post-deafening changes in neuronal and neurotrophic factor gene expression

Death of SGNs after hair cell loss is gradual, with 80%-90% of SGNs lost over a period of at least 14 weeks in aminoglycoside-treated rats (Alam et al., 2007). However, there are functional and morphological changes in deafferented SGNs even while they are still alive (Shepherd and Hardie, 2001). SGNs surviving after kanamycin-induced hair cell loss in guinea pigs showed shrinkage of the neuronal soma and peripheral axons (Wise et al., 2017). This axonal and somatic atrophy may be related to reduced expression of genes encoding cytoskeletal elements, including neurofilaments (Nefh, Nefm, Nefl) and microtubules (Tubb3, Map1a). However, the reduced gene expression is not evident until P60 in our dataset and the atrophy reported by Wise et al. (2017) is already evident at two weeks post-deafening. The reduced gene expression may simply be due to the reduced number of neurons at P60.

Functional changes in SGNs include lengthened action potential refractory period and elevated threshold observed in auditory nerve recordings from aminoglycoside-deafened rats (Shepherd et al., 2004). Consistent with this, we observed downregulation of K_v_3 genes (Kcnc1 and Kcnc3) necessary for the high-frequency firing rates (Rudy and McBain, 2001; Lien and Jonas, 2003) that are characteristic of SGNs. Significant changes can also be seen at the auditory nerve synaptic terminals on spherical bushy cells in the cochlear nucleus of aminoglycoside-deafened or congenitally deaf cats, including >35% reduction in the number of synaptic vesicles (Ryugo et al., 1997; Ryugo et al., 2010; O’Neil et al., 2011). Consistent with these findings, we observed downregulation of several genes involved in presynaptic functions, particularly those associated with synaptic vesicle trafficking and transport (Figure 4D). It could be argued that this is simply a result of the fact that SGNs are dying so all neuronal gene transcripts are decreasing. However, downregulation of genes related to neuronal channels and synaptic function is already occurring at P32, prior to the onset of SGN death and a time at which abundant neuronal transcripts such as those related to cytoskeletal elements are not significantly downregulated.

Interestingly, among neuronal genes *not* downregulated post-deafening are genes encoding NTFs and their receptors. We have previously shown (Bailey and Green, 2014) that, after deafening, NTFs continue to be expressed in the the organ of Corti and cochlear nucleus, respectively, the pre-and postsynaptic targets of spiral ganglion neurons. Here, we show that this is also the case for NTFs synthesized in the ganglion. These results showing continued availability of NTFs post-deafening support the hypothesis that SGN death is not a consequence of loss of neurotrophic support.

### Post-deafening changes in Schwann cells

SGNs are not the only cells to undergo changes after deafening. Spiral ganglion Schwann cells also alter their morphology and phenotype and, presumably, this is reflected in some of the transcriptional changes in our RNAseq dataset. Schwann cells have been shown to convert to a non-myelinating phenotype (Wise et al., 2017), accompanied by increased expression of the NTF receptor p75^NTR^ (Provenzano et al., 2011). Also, Schwann cells enter cell cycle and proliferate postdeafening (Provenzano et al., 2011). Correspondingly, we observed increased expression of the p75^NTR^ gene (Ngfr) in P60D vs. P60H as well as an enrichment of GO categories for cell division and cell cycle (Figure 3B) and glial proliferation/activation (Figure 2). The reasons for these changes and the functions of these newly dedifferentiated Schwann cells remains unclear, but it has been suggested that proliferation and conversion to a non-myelinating phenotype would allow Schwann cells to support deafferented SGNs and promote neural regrowth (Provenzano et al., 2011; Wise et al., 2017), which may include phagocytosis of myelin and neuronal debris. Another interesting possibility is that the dedifferentiated Schwann cells are participating in recruitment of macrophages (Nazareth et al., 2021) and triggering the post-deafening immune response.

### Are macrophages detrimental or beneficial to SGN survival?

Macrophages are present in the cochlea of normal hearing mice (Hirose et al., 2005; Kaur et al., 2015) and humans (Liu et al., 2018). Here we confirm our previous observations (Rahman et al., 2023) that macrophages are present in the spiral ganglion of normal hearing rats. Following acoustic or ototoxic trauma in mice (Kaur et al., 2018; Bedeir et al., 2022) or rats (Rahman et al., 2023), macrophage numbers in the spiral ganglion significantly increase as SGN numbers decrease. Here, we similarly observed increased macrophage density in the spiral ganglion of aminoglycoside-deafened male and female rats at a time when SGNs are dying (Figure 9B, D). Although there is temporal correlation between SGN death and the increase in macrophage number, there is a lack of spatial correlation. Macrophage density is highest in the apical cochlea while the decline in SGN density appears to be greatest in the middle of the cochlea (Figure 7D,E). Thus, if macrophages play some role in SGN death, direct contact between macrophages and SGNs may not be necessary.

It has been suggested that macrophages play a neuroprotective role in the spiral ganglion; disruption of fractalkine signaling, one of the chemoattractants for macrophages, results in reduced macrophage infiltration and decreased SGN survival after cochlear trauma (Kaur et al., 2015; Kaur et al., 2018; Stothert and Kaur, 2021). However, in this study, macrophages, while impaired in guided mobility, remained present in the spiral ganglion and other studies have suggested that macrophages may promote SGN death. Anti-inflammatory agents that reduced macrophage numbers or activation in the spiral ganglion, had the effect of reducing SGN death post-deafening (Rahman et al., 2023) and depletion of macrophages by systemic injection of clodronate, also reduced SGN death postdeafening (Shimada et al., 2023). Moreover, systemic lipopolysaccharide, a potent macrophage chemoattractant and activator, exacerbated kanamycin and furosemide ototoxicity (Hirose et al., 2014). Together, these results suggest that, while macrophages contribute significantly to SGN death postdeafening, some macrophages may be neuroprotective. This is likely a consequence of the heterogeneity of the cochlear macrophage population following ototoxic trauma (Bedeir et al., 2022). Similarly, in other studies of macrophage involvement in trauma in the CNS (Chen et al., 2015) or non-infectious diseases generally (Yousaf et al., 2023), different macrophage types – e.g., M1 and M2 – can promote or reduce inflammation with consequent neurotoxic or neuroprotective consequences.

While inflammatory and immune responses to cochlear damage have been well documented, of the several cues available for their recruitment (Warchol, 2019), it is not yet clear which is/are directly responsible. We hypothesized that bacteria or other pathogens can gain access to the cochlea after trauma, perhaps due to disruption of the blood-labyrinth barrier, and constitute the proximate trigger for neuroinflammation. For example, it has been shown that bacteria are present in the brain of Alzheimer’s disease patients (Emery et al., 2017) and the infection by bacteria and/or viruses may contribute to the pathogenesis of Alzheimer’s disease (Piekut et al., 2022). We used BLAST to assess whether any of the RNAseq reads that did not map to the rat genome had significant alignments to bacterial sequences. We found that around 12-15% of unmapped reads had at least one significant alignment to a bacterial sequence. However, these did not differ significantly between hearing and deafened samples, making it unlikely that bacteria are causing the post-deafening immune response.

### Possible mechanisms for innate immune-mediated SGN death

Innate immune response genes exhibited increased expression after deafening. These include several inflammasome genes, including Nlrp3, Pycard, Casp1, and Aim2. This implicates, in SGN degeneration, inflammasome activation, which has been shown to be involved in genetic (Nakanishi et al., 2020) and cytomegalovirus-induced (Zhuang et al., 2018) hearing losses. One possible mechanism is induction of inflammasome activation by ATP (Oliveira-Giacomelli et al., 2021), which is released by cochlear sensory epithelial cells after trauma (Lahne and Gale, 2008). While there are many possible mechanisms by which the innate immune response could be causing SGN death, here we focus on two, the complement system and recruitment of the adaptive immune system.

#### Complement

Complement genes were among the immune response-related genes upregulated after deafening, indicating activation of the complement pathway in the spiral ganglion post-deafening. Complement activation has been shown to promote neuronal pathologies in the brain; including those associated with traumatic brain injury (Alawieh et al., 2021), Alzheimer’s disease (Fonseca et al., 2004), multiple sclerosis (Morgan et al., 2020); and peripheral neuropathies (Dailey et al., 1998; Xu et al., 2018; Royer et al., 2019). Nevertheless, we found that deletion of C3, a key component in complement activation, did not improve SGN survival (Figure 8) nor reduce macrophage infiltration (Figure 8) or activation (Supplemental Figure 6).

This suggests that complement is not required for immune response activation post-deafening nor for subsequent SGN degeneration. However, there are other possible roles for complement in the response to cochlear trauma. Complement activation can be involved in promoting clearance of cellular debris and pathogens (Noris and Remuzzi, 2013), and complement is one of the mechanisms linking innate to adaptive immune responses (Royer et al., 2019; Lo and Woodruff, 2020).

#### Adaptive immune responses

Genes associated with the adaptive immune response were among the genes upregulated after deafening. This was supported by our observation of significant increases in the number of lymphocytes and MHCII-expressing macrophages in the spiral ganglion after hair cell loss (Figure 11). Moreover, the presence of cells of the innate and adaptive immune systems in the cochlea and their increase after noise damage has been previously reported (Rai et al., 2020). Together, these results indicate recruitment of both the innate and adaptive immune systems after cochlear trauma. It has been shown that T cells are involved in CNS neurodegenerative diseases (Subbarayan et al., 2020) and peripheral neuropathies (Royer et al., 2019). Further studies will be needed to determine the extent to which the innate and adaptive responses occurring post-deafening contribute to SGN death.

Maintaining SGN survival and physiological function is important for cochlear implant patients. Our observations that both innate and adaptive immune responses are recruited after hair cell loss and may play a role in SGN degeneration or death, have implications for development of therapeutic strategies. However, effectively targeting the relevant components of the complex immune response requires clarification of the precise mechanisms utilized by the immune system to prevent or cause SGN death postdeafening.

## Abbreviations

ABR: auditory brainstem response
C3: complement component
C3: GO Gene Ontology
GSEA: Gene set enrichment analysis
IL: Interleukin
KO: knockout
log_2_FC: log_2_ fold-change
padj: adjusted p-value **P** postnatal day
P0: postnatal day 0 (birth)
P32D: postnatal day 32 deafened
P32H: postnatal day 32 hearing control
P60D: postnatal day 60 deafened
P60H: postnatal day 60 hearing control
PBS: phosphate buffered saline, pH 7.4
PCA: principal component analysis
PFA: paraformaldehyde
qPCR: quantitative real-time polymerase chain reaction
SGN: spiral ganglion neuron
TBS: Tris-buffered saline, pH 7.6
WT: wild-type

## Acknowledgements

We thank Catherine Kane for maintaining our animal colony and her assistance in collecting cochlear tissue for the RNAseq experiments. We also thank the Iowa Institute for Human Genetics Genomics Division for performing the library preparation and RNA sequencing, Suchita Komanduri for assistance in histological processing and labeling of cochlear sections, and members of the Green lab for valuable feedback on the manuscript.

## Scope statement

This manuscript provides novel information on spiral ganglion neurodegeneration after ototoxic trauma – hair cell death due to aminoglycoside ototoxicity. RNAseq was used to document transcriptional changes occurring in the spiral ganglion of the cochlea after deafening. We identify genes transcriptionally *down*regulated after deafening that are responsible for degradation of specific neuronal functions important for neurotransmission by the auditory nerve. We also show that the largest group of transcriptionally *up*regulated genes are for immune responses. These involve recruitment of the complement pathway, innate immune responses, and adaptive immune responses (although we show that the complement pathway is not necessary for spiral ganglion neuron death). We confirm these results by histology to show that the spiral ganglion post-deafening has increased numbers of cells of the innate and adaptive immune systems. Thus, we show that the cochlear response to ototoxic trauma includes inflammatory, innate immune, and adaptive immune components associated with degeneration and death of spiral ganglion neurons.

## Data availability

The accession number for the data reported in this paper is GSE194063, which contains the raw fastq files and matrices of raw and normalized read counts. Code/scripts and other analysis files not already made available will be made available from authors upon reasonable request.

## Supplementary figure legends

**Supplementary Figure 1.** Experimental procedures. A) Schematic depicting the dissection procedure for collecting tissue for RNA. The cochlea was isolated from the temporal bone and the bony capsule removed. The organ of Corti was then removed, leaving the spiral ganglion and associated cells. Cochleae were collected and RNA extraction performed as described in Materials and Methods. B) Representative midmodiolar section of the cochlea. The different half turns are outline and labeled in green. C) Western blot of liver protein lysates to detecting C3 protein in WT and heterozygous rats and confirming lack of C3 protein in C3KO rats. KM: kanamycin; ABR: auditory brainstem response; SGN: spiral ganglion neuron.

**Supplementary Figure 2.** Principal Component Analysis of RNAseq replicates. A) PCA plot of all biological replicates. Note than one P32H replicate does not cluster with the other P32H replicates. B) PCA plot after the variant P32H replicate was removed. Replicates for each condition are clustered similarly.

**Supplementary Figure 3.** Number of categories in each cluster from GSEA network visualizations. A) The number of categories in each cluster of the overall hearing vs. deaf network (Fig. 1C). B) The number of categories in each cluster of the P32 hearing vs. deaf network (Fig. 2A). C) The number of categories in each cluster of the P60 hearing vs. deaf network (Fig. 2B). The intracellular signaling group is subdivided into two clusters; the immune response group is subdivided into 10-12 clusters with the numbers in the legends corresponding to the numbered clusters in each respective Fig. 1C or Fig. 2 network.

**Supplementary Figure 4.** C3KO does not affect normal hearing. A) Representative waveforms from hearing WT (black), heterozygous (blue), and C3KO (orange) rats elicited by suprathreshold stimuli at 8 kHz. B) Graph showing ABR thresholds for 4 kHz, 8 kHz, and 16 kHz tone pips in WT, heterozygous, and C3KO rats. C-D) Graphs showing wave I amplitude (C) and wave I latency (D) for 4 kHz (left), 8 kHz (middle), and 16 kHz (right). Data were fit to 2^nd^ order polynomial curves (lines). WT: wildtype; het: heterozygous; KO: knockout. n = number of ears.

**Supplementary Figure 5.** Kanamycin-induced hair cell loss is not different in C3KO rats. A-C) Representative images of cochlear wholemounts labeled for hair cells (Myo6/7a, cyan) and neurons (NF200, magenta) from hearing WT (A), heterozygous (B), and KO (C) rats. Hair cells appeared normal in all three genotypes. D-F) Representative images of cochlear wholemounts labeled for hair cells and neurons from kanamycin-injected WT (D), heterozygous (E), and KO (F) rats. Hair cell death was similarly essentially complete in all three genotypes.

