## Supplementary Figures and Table for "Spiral ganglion neuron degeneration in deafened rats involves innate and adaptive immune responses not requiring complement"

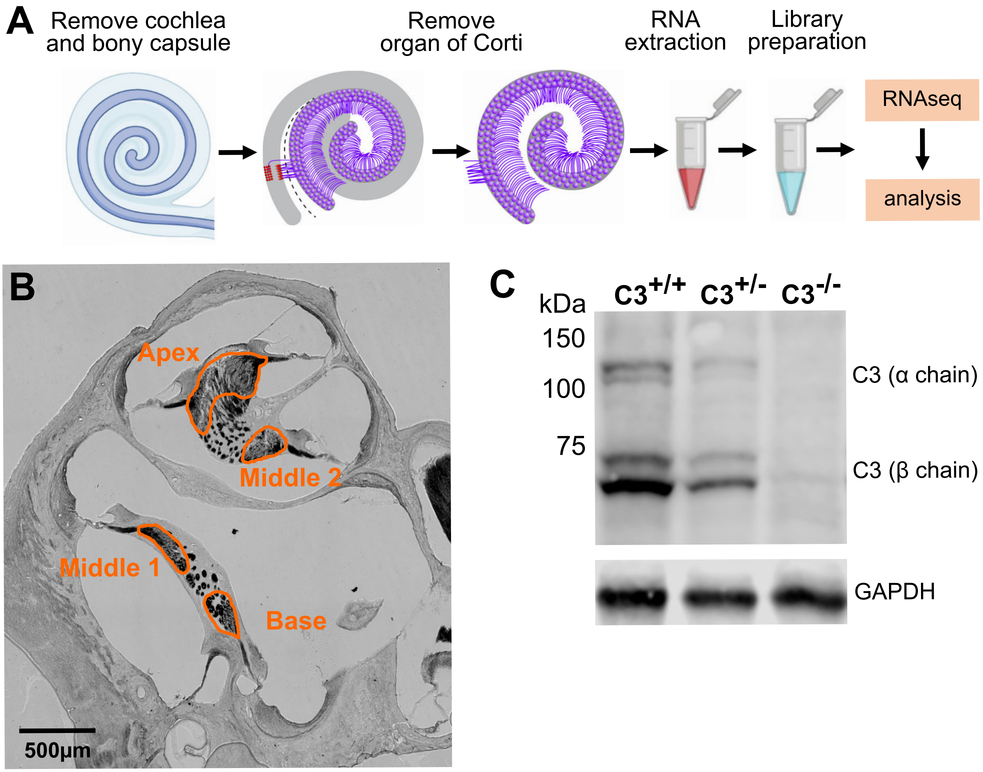


Supplementary Figure 1

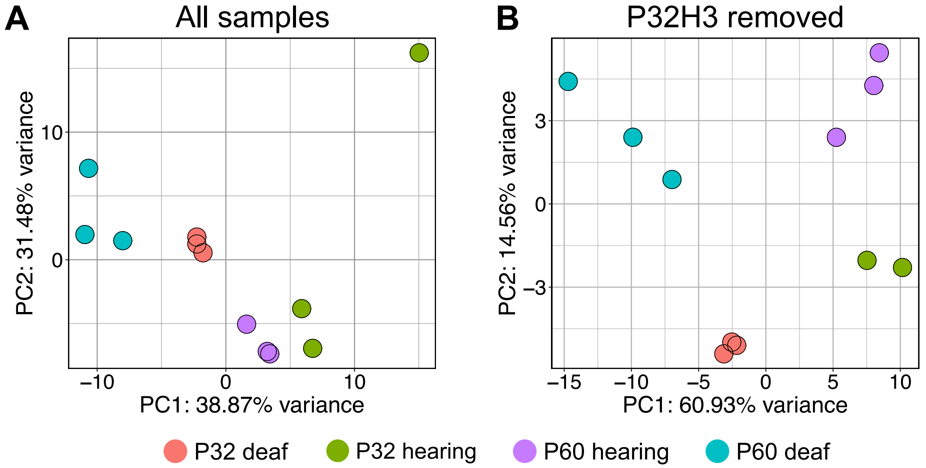


Supplementary Figure 2

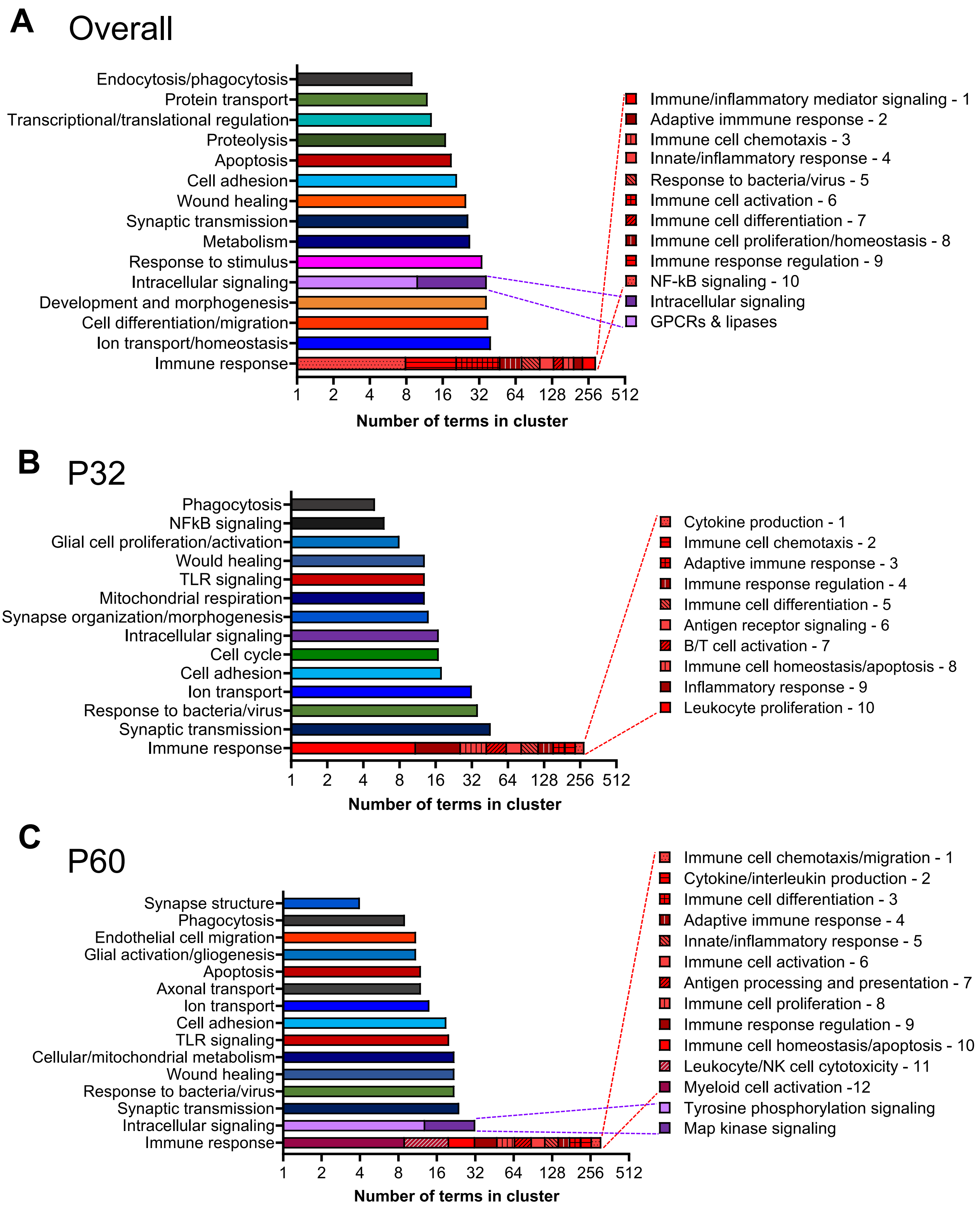


Supplementary Figure 3

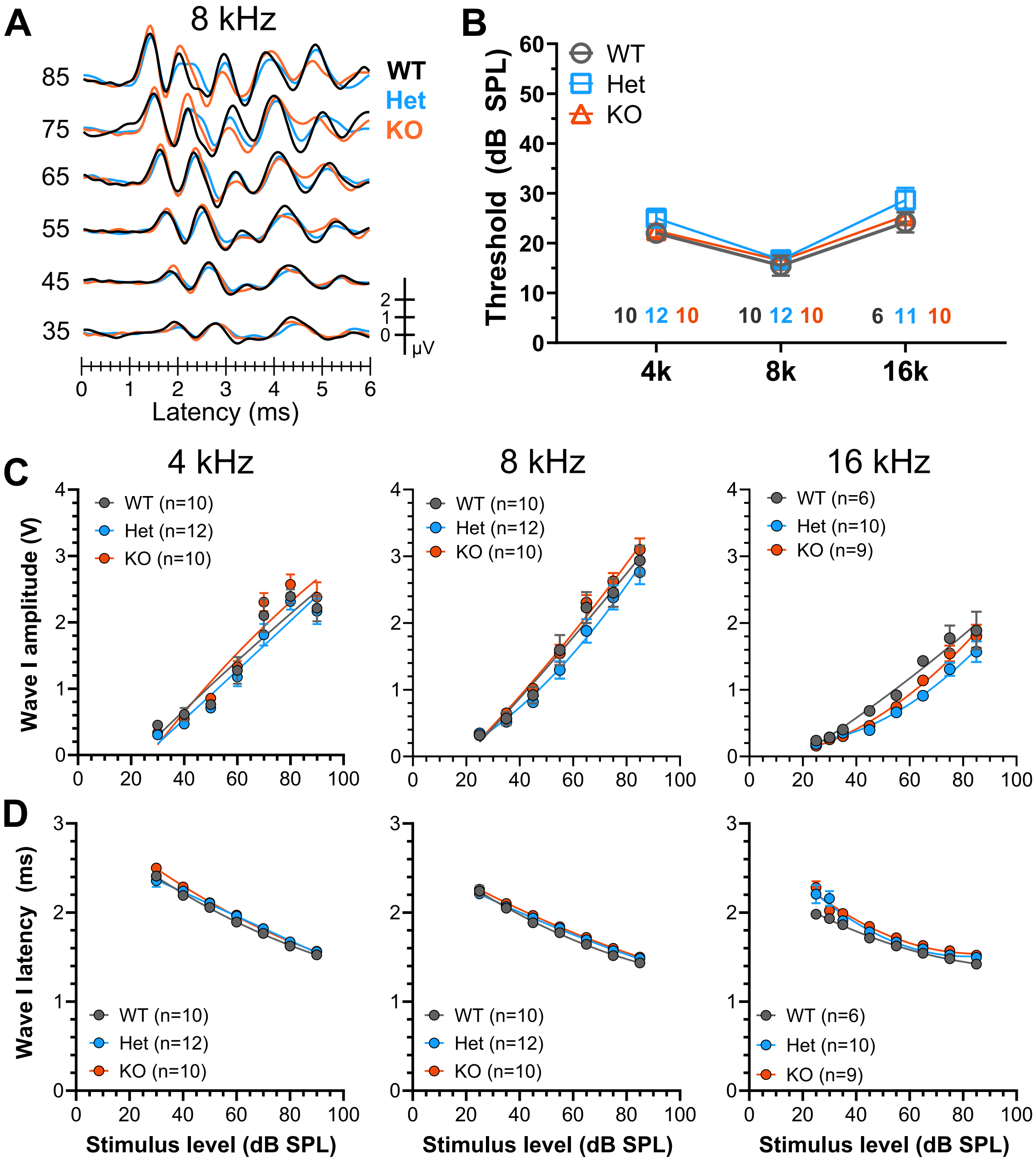


Supplementary Figure 4

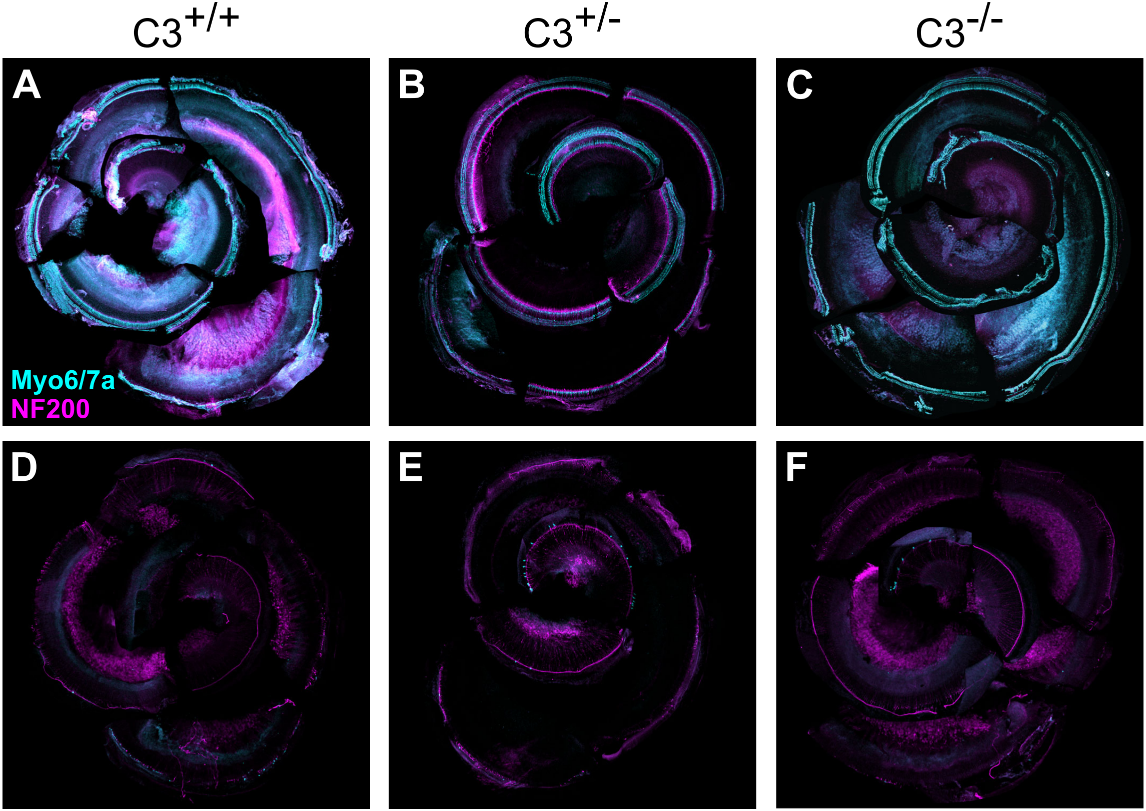


**Supplementary Figure 5. Cochlear wholemounts labeled to show hair cells and neurons.** Hair cells are labeled with Myo6/7a, neurons with NF200 antibodies. Genotypes are shown above each column: **A&D**, wildtype; **B&E**, heterozygotes; **C&F**, homozygous C3 knockouts. **A-C**, control undeafened; **D-F**, deafened with kanamycin.

Supplementary Figure 5

**Supplemental Table 1. qPCR primers**

| Gene name | Primer sequences | Accession ID | Amplicon length (bp) |
|---|---|---|---|
| C1qa | F: CCGGGTCTCAAAGGAGAGAG<br>R: CCAGATTCCCCCATGTCTC | NM_001008515.1 | 89 |
| C3 | F: CAGTGTGGGTGGATGTGAAG<br>R: GGCTGTCGGTTATCTCTTGG | NM_016994.2 | 76 |
| Cd4 | F: CCCTGAACCAGAAGAAGCAC<br>R: CAAGGTTGAGTGGGAAGGAG | NM_012705.1 | 123 |
| Cd68 | F: CTCATTCCCTTACGGACAGC<br>R: GCTGAGAATGTCCACTGTGC | NM_001031638.1 | 135 |
| Cxcl10 | F: GCTTATTGAAAGCGGTGAGC<br>R: GGGTAAAGGGAGGTGGAGAG | NM_139089.1 | 122 |
| RT1-Da | F: AAGAAAATGGCCACACTTGG<br>R: CGGGGGAAAGATAGAACTCC | NM_001008847.2 | 130 |
| Kcnc1 | F: CCTACTCATCCCGCTATGC<br>R: GTGGGTTCTCAGACAGAAGG | NM_012856.2 | 89 |
| Kcnc3 | F: ACTGGGAATTATGGGATTGC<br>R: AGTCCTCATGAGCCAGTGC | NM_053997.5 | 140 |
| Kcnd2 | F: AGCCTTCTGGTACACCATCG<br>R: TTCCCTGCTATGGTTTTTGG | NM_031730.2 | 75 |
| Nefh | F: CCGTCATCAGGTAGACATGG<br>R: AGCAGGTCCTGGTACTCTCG | NM_012607.2 | 117 |
| Scn4b | F: GAACCGAGGCAAATACTCAGG<br>R: CAGATACCTCCAACGACAGG | NM_001008880.2 | 89 |
| Scn8a | F: AGCGCTGAGAACATTCAGG<br>R: CTTCACGGACTGGATTAGGG | NM_019266.3 | 97 |
| Syp | F: CCAGACAGGGAACACATGC<br>R: AGCCTGTCTCCTTGAACACG | NM_012664.3 | 131 |
| Rps16 | F: GAAGGGTGGTGGTCATGTG<br>R: TCCGATCGTACTGGATGAGG | NM_001169146.1 | 137 |
